# The *B*-value calculator: expected diversity with background selection

**DOI:** 10.64898/2026.03.04.709642

**Authors:** Jacob I. Marsh, Austin T. Daigle, Parul Johri

## Abstract

Background selection (BGS), the indirect effect of purifying selection at linked and unlinked sites, is a key evolutionary process shaping genomic patterns of variation. Calculating the expected diversity at neutral sites experiencing BGS relative to that under strict neutrality (referred to as *B*-value or simply *B*) is important for developing null models when performing population genomic inference, in particular, demographic inference and detection of selective sweeps. We extend and integrate previous theory to estimate *B*-values analytically, assuming no selective interference, with novel expressions to account for gene conversion between proximal sites. Here, we present the *B*-value calculator, *Bvalcalc*, an easy-to-use command-line interface written in Python for efficient analytical calculation of expected genome-wide *B* at single base-pair resolution. *Bvalcalc* has several modules for calculating diversity as a function of distance from a single selected element or considering the multiplicative effects of all selected elements across the genome, accounting for recombination maps, gene conversion, self-fertilization, single population size changes, and unlinked effects from other chromosomes. We validated the effectiveness of *Bvalcalc* with comparisons against simulated results, and generated *B*-maps for the model species *Homo sapiens*, *Drosophila melanogaster* and *Arabidopsis thaliana* as a proof of concept using public data. *Bvalcalc* is available with documentation at johrilab.github.io/Bvalcalc/.

**Author Summary:** Selection against deleterious alleles reduces diversity and distorts the site frequency spectrum at linked loci and across the genome. This phenomenon is termed background selection (BGS). BGS can confound population genetic analyses that assume strict neutrality, including inferring historical population size changes or scanning for selective sweeps from beneficial mutations. Here, we present *Bvalcalc* an easy-to-use command-line tool for calculating the expected diversity with BGS (*B*) relative to that under neutrality, while accounting for genome architecture. We extend and integrate many analytical expressions into a single accessible tool so that users may provide population-specific parameters such as effective population size, crossover rate, mutation rate, selfing rate, and a distribution of fitness effects (DFE) and calculate the expected BGS effects from genomic elements experiencing direct purifying selection. We validate the effectiveness of *Bvalcalc* against simulated results and find that *Bvalcalc* can accurately estimate the effects of BGS under most scenarios. As a proof of concept, we build *B-*value maps for *Homo sapiens*, *Drosophila melanogaster* and *Arabidopsis thaliana*. *Bvalcalc* is designed to be applicable to non-model species.

## Introduction

Background selection (BGS) refers to the indirect effects of selection at linked and unlinked sites due to directly selected sites experiencing purifying selection. BGS is an important evolutionary process that alters evolutionary dynamics for sites proximal to selected elements and across the genome [1]. The effect of BGS on diversity is quantified by the ratio of diversity at a given neutral site embedded in a genome experiencing purifying selection (*π*), compared to expected diversity (*π*_0_) under identical conditions in a strictly neutral population. A *B*-value is defined as *B* = *π*/*π*_0_.

Besides reducing the effective population size (*N_e_*) [2], BGS confounds demographic inference by distorting the site frequency spectrum of linked variants (SFS)[3,4], and local genealogies in the ancestral recombination graph (ARG)[5]. BGS can significantly decrease nucleotide diversity in regions of low recombination, which may lead to falsely characterizing those regions as selective sweeps ([6,7], although see [8]). Additionally, researchers need to account for the expected effects of BGS to understand the contribution of recurrent selective sweeps to shaping levels of genomic variation [9–11]. Quantifying the effects of BGS by accurately calculating *B*-values across the genome (generating *B*-maps) is therefore essential for developing an effective null model for population genetic inference [12–15].

Classic mathematical models of background selection [1,2,16,17] enable researchers to mathematically predict the effect imposed by a selected site experiencing purifying selection on nucleotide diversity at a linked site. Population geneticists have evaluated and developed expressions to account for how *B*-values are affected by the presence of gene conversion [18,19], self-fertilization (selfing)[20–24], unlinked BGS [25,26], distributions of fitness effects for selected loci (DFE; *e.g.*, [14,27]), non-equilibrium demography [4,28–31]. Additional expressions have been developed for higher-order Hill-Robertson interference effects (HRI) between co-segregating alleles experiencing purifying selection [25,32–36,76]. However, there is a pressing need for effective integration of these advances into unified models applicable to real genomes, so that population genetic theory can be readily applied to empirical problems.

Algorithms implementing BGS models to genomes with realistic architecture have allowed for the calculation of *B*-maps in select species, most notably humans [27,35,37] and *Drosophila melanogaster* [10,12]. These *B*-maps have been generally effective at explaining observed diversity patterns across the genome. *B*-maps constructed with these algorithms have been applied to assess and mitigate the effects of BGS on demographic inference [38,39], improve reliable selective sweep detection [6,40], identify biases in fine-scale recombination rate inference [41], and improve GWAS [42]. However, *B*-maps are scarcely available beyond these model systems and the publicly available software tools are foundational and effective, but can be difficult to approach for all but those with highly technical expertise.

Here, we present the *B*-value calculator, *Bvalcalc*, an accessible and flexible Python package that enables users to calculate accurate *B*-values analytically at single base pair resolution for model and non-model species. We have incorporated several key extensions of the classic BGS model from the literature, validated with simulated results. Features include integration over a DFE, BGS effects from unlinked sites, support for selfing populations, recombination rate maps, and simple demographic changes in a two-epoch model. In addition, we provide extended expressions accounting for gene conversion. Note that *Bvalcalc* does not model HRI by default. We intend *Bvalcalc* to be of broad utility to technical and non-technical population biologists, and so have developed comprehensive documentation with guides, example code snippets, and other learning materials to aid its accessibility (see https://johrilab.github.io/Bvalcalc/). To this end, we also provide several pre-built templates of the population genetic parameters (*i.e.,* parameters that may be used to effectively predict *B*-values from human coding sequences) that can be retrieved from the *Bvalcalc* command-line interface. Finally, we use *Bvalcalc* to calculate *B*-maps for the model species human, *D. melanogaster* and *Arabidopsis thaliana* as a proof of concept and as resources for the community (available at https://johrilab.github.io/Bvalcalc/prebuilt_maps/bmaps.html).

## Methods

### Obtaining *B*-value

*Bvalcalc* calculates the BGS effect imposed by a functionally important genomic element *i* experiencing purifying selection, on a focal site at an arbitrary distance (*y*) from the end of the element. Assuming that there are *E* functional genomic elements on a chromosome, the final *B*-value (*B_tot_*) at the focal site is given by the product of the BGS effects caused by all selected elements on the focal chromosome and unlinked effects (*B_unlinked_*) from other chromosomes, as illustrated in Figure 1. That is,

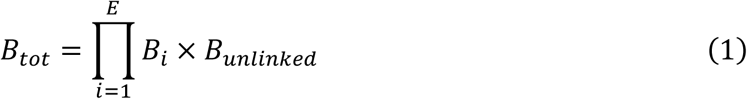

Let *ϕ*(*t*) be the probability density function of *t* (= *sh*), where *s* is the selective effect of a homozygous mutant relative to wild type and *h* is the dominance coefficient. We assume that *ϕ*(*t*) is given by the combination of four non-overlapping uniform distributions denoting the effectively neutral class (0 ≥ 2*N_anc_s* > −1), weakly deleterious class (−1 ≥ 2*N_anc_s* > −10), moderately deleterious (−10 ≥ 2*N_anc_s* > −100) class and strongly deleterious (−100 ≥ 2*N_anc_s* > −2*N_anc_*) class of mutations, such that they are present in proportions *f*_0_, *f*_1_, *f*_2_, and *f*_3_, respectively (i.e., ∑*_i_ f_i_* = 1). Here, *N_anc_* refers to the ancestral effective population size. Let *t_i_* (= -*s*_i_*h*) represent the boundary of these uniform distributions, scaled by *h*; e.g., *t*_2_ = *h* × (10 / (2 × *N_anc_*)). Here we assume that the rate of crossover per site/generation is denoted by *r*, the rate of gene conversion per site/generation is denoted by *g* and the gene conversion tract length (*k*) is fixed. The total deleterious mutation rate across an element is equal to *U* = 2*μL*, where *μ* is the mutation rate per site/generation. For any genomic element of length *L* that is at distance *y* from the end of the element, we integrate over the length of the element and the DFE, *ϕ*(*t*), as done in Johri et. al (2020)[14]. Let us write *B_i_* = *exp*(−*H*). Then we obtain,

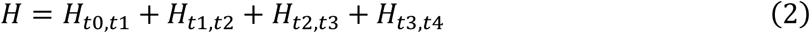

where

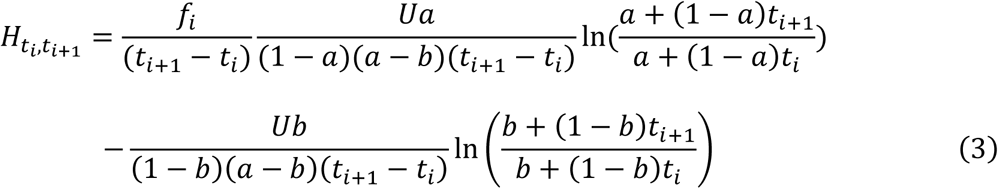

where *a* = *gy* + *C* and *b* = *g*(*y* + *L*) + *rL* + *C* when *y* + *L* << *k*; and *a* = *gk* + *C* and *b* = *gk* + *rL* + *C*, otherwise. Here, *H_ti_*_,*t*_*_i_*_+1_represents the contribution of BGS from deleterious mutations that belong to the uniform distribution given by bounds *t_i_* (= −*hs_i_*) and *t_i_*_+1_ (=−*hs_i_*_+1_). Note that while Equation 2 here assumes four uniform distributions, in the method, the user has the option to implement any arbitrary number of distributions. It should also be noted that these derivations assume that 2*N_anc_ sh* ≪ −1. We therefore ignore the contributions of weakly deleterious mutations (*i.e*., those with 2*N_anc_sh* > −5) to BGS, which has been previously shown to approximate the effects of BGS well even when the DFE comprises predominantly weakly deleterious mutations (see Figure 1D of Johri et al. 2020[14]).

Here,

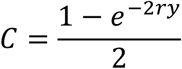

The parameter *C* accounts for multiple crossovers by employing the Haldane function [43], which assumes that there is no crossover interference. The derivation of the above equation is shown in Supplemental Text S1. Note that the terms *a* and *b*, which capture the crossover and gene conversion effects, were further modified to account for complexities involved in short-range linkage under gene conversion. These new expressions are presented in Supplemental Text S2.

**Figure 1:**
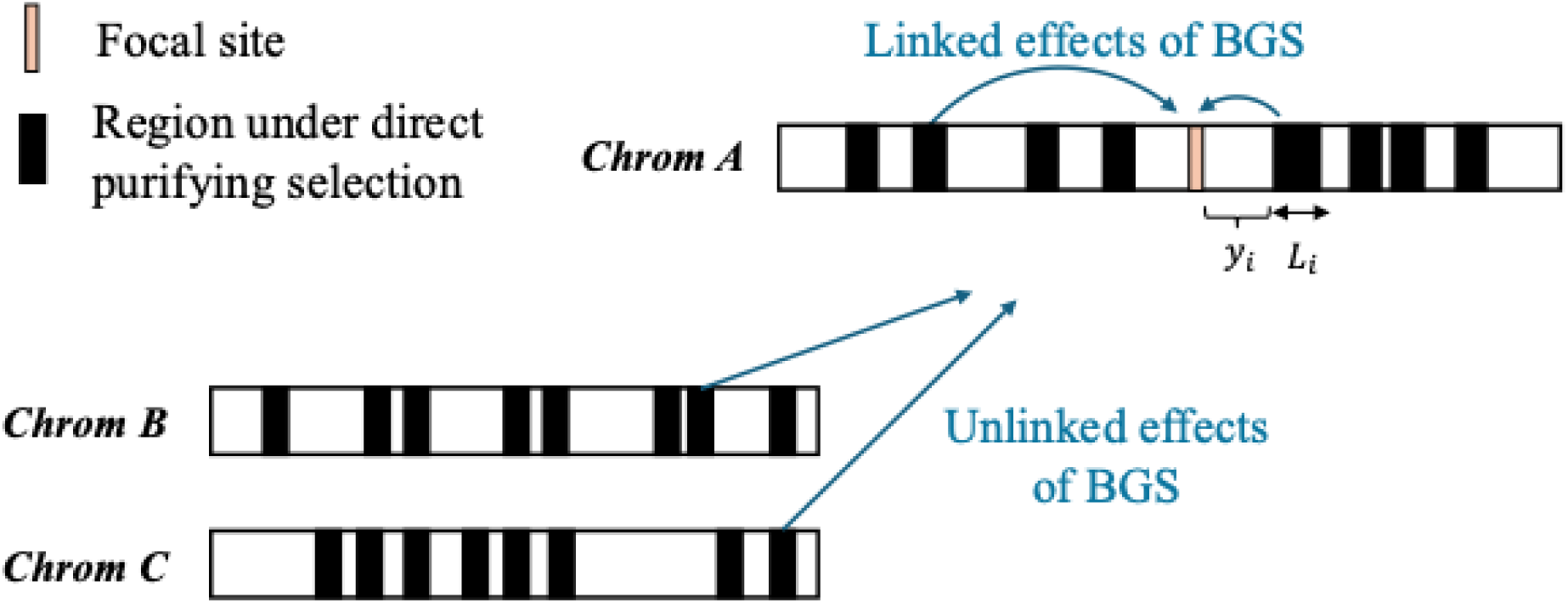
Illustrative diagram of the combined effects of background selection (BGS) at a focal site from linked and unlinked regions experiencing direct purifying selection (selected elements). All selected elements contribute to BGS, reducing diversity at a given focal site, which is accounted for by *Bvalcalc* when calculating *B-*values across the genome. *L*_i_ represents the length of the *i*^th^ selected element and *y*_i_ is the distance of the focal site from the end of the same selected element.

#### Effects of selfing

Self-fertilization was modelled by modifying the relevant population genetic parameters by Wright’s inbreeding coefficient (*F*; [44]), which is a function of the selfing rate (*S*) such that *F* = *S*/(2 − *S*) [21]. Our implementation includes a reduction of the population size to an effective selfing-adjusted population size, *N_eff_* = *N* / (1 + *F*), a decrease in the crossover rate, *r_eff_* = *r* × (1 − *F*), a similar decrease in the gene conversion initiation rate, *g_eff_* = *g* × (1 − *F*) and a modified dominance coefficient, *h_eff_* = *h* + *F*(1 − *h*).

#### Unlinked effects of BGS

Let us assume that there are *M* chromosomes and *L_j_* is the length of chromosome *j*. Let us also assume for the sake of simplicity that *f_sel_* represents the average proportion of selected sites on all chromosomes. Then, unlinked effects of BGS at the focal site (on chromosome *j*) would be given by

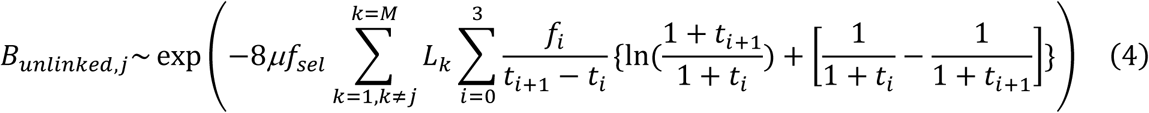

See the Supplemental Text S1 for the derivation of analytical expressions to obtain the unlinked effects of BGS. Equation 4 is consistent with expressions derived assuming free recombination [2,23].

#### Accounting for a single instantaneous size change

We assume that the population changes in size from *N_anc_* to *N_cur_* at time *T* (generations) in the past, and that *B_tot_*_,*anc*_ and *B_tot_*_,*cur*_ are the values of *B_tot_* in the ancestral and current population, respectively. Then using Equation 1b from Johri *et al.* (2021)[4], *B_tot_*_,*cur*_ can be obtained as:

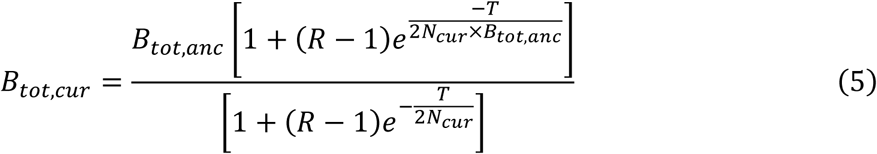

Here *R* = *N_anc_* ⁄*N_cur_*. Note that these derivations assume that *T* is small enough that *B_tot_*_,*cur*_ is more or less constant across time 0 to *T*.

#### Accounting for Hill-Robertson interference (HRI)

In this study, we define *B*-values assuming no HRI effects. *B*-values accounting for the presence of HRI, referred to as *B*’-value 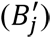 are not supported in *Bvalcalc* by default. See Text S4 for an extension of Becher and Charlesworth (2025)[36] to calculate 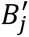 for a non-recombining region in diploids and to account for a DFE using the coarse-graining method described by Good *et al.* (2014)[33].

For software details regarding the algorithms implemented in *Bvalcalc* to calculate *B*-maps, see Text S3.

### Simulations and the basic model

A diploid Wright-Fisher model was simulated using *SLiM* 4.0.1 [45]. The basic model included population parameters resembling those of *D. melanogaster*, albeit with an arbitrary deleterious DFE (see below) and a simple genomic architecture. An effective population size of *N* = 1 × 10^6^ was assumed, with a constant mutation rate of *μ* = 3 × 10^−9^ per site/generation [46]. A relatively low sex-averaged constant cross-over rate of *r* = 0.5 × 10^−8^ per site/generation [47] was assumed to observe stronger effects of BGS. A selected element of 10 kb in length was simulated with a 40 kb linked neutral region on one side for testing our extensions of the classic BGS model. When testing the effects of multiple selected genomic elements, 10 elements each of length *L* = 10 kb were spaced 10 kb apart in a 200 kb genome. The genomic elements experienced purifying selection such that the DFE was comprised of four non-overlapping uniform distributions representing effectively neutral mutations (−1 ≤ 2*Ns* < 0), mildly deleterious mutations (−10 ≤ 2*Ns* < −1), moderately deleterious mutations (−100 ≤ 2*Ns* ≤ −10), and strongly deleterious mutations (−100 ≤ 2*Ns*) present in proportions *f*_0_, *f*_1_, *f*_2_, and *f*_3_, respectively. The DFE is defined by the frequency of each class of mutation at a directly selected site such that *f*_0_ + *f*_1_ + *f*_2_ + *f*_3_ = 1. All mutations were assumed to be semidominant unless specified otherwise.

For the basic model, the DFE was *f*_0_ = 0.1, *f*_1_ = 0.2, *f*_2_ = 0.3, *f*_3_ = 0.4. Simulations were rescaled by a factor of 100 such that the simulated population size was *N_scaled_* = 10^4^, following recommendations provided by Marsh *et al.* (2026)[48]. Mutation rates and recombination rates (crossover and gene conversion) as well as selection coefficients were multiplied by the scaling factor (100). For all results, the population was simulated with a burn-in of 14*N_anc_* generations, where *N_anc_* is the ancestral population size. 100 diploid genomes were sampled at the end of the burn-in and nucleotide diversity at neutral sites was calculated across 10000 replicates unless otherwise stated. To ensure that 14*N_anc_* generations were sufficient to reach equilibrium levels of diversity, we compared mean diversity at neutral sites obtained from simulations with a burn-in period of 13*N_anc_* vs 14*N_anc_* generations. A 40kb neutral region was simulated linked to a 10kb selected element under the basic model. We observed no significant differences in mean *B* values when averaged across 10000 replicates (Figure S1, Figure S2).

The following extensions were added to the basic model: A gene conversion rate of *g* = 1 × 10^−8^ per site/generation with a geometrically distributed tract length of mean 440 bp, with only simple conversions (not involving heteroduplex mismatch repair). Instantaneous population size expansion (*N_cur_* = *N_anc_* × 5) as well as instantaneous contraction (*N_cur_* = *N_anc_* / 5), where *N_cur_* is the current population size since the change, that occurred after the burn-in (14*N_anc_* generations) in simulations with demography. For these simulations we sampled genomes post-change corresponding to *T* = 0.2 and *T* = 1, *T* = 2, where *T* is measured in units of *N_anc_* generations. For instance, when the population underwent a five-fold growth from *N_anc_* = 10^4^ to a current population size of size *N_cur_* = 5 × 10^4^, sampling was performed at 2,000 generations post-change (*T* = 0.2), and 10,000 generations post-change (*T* = 1). A selfing rate (*S*) of 0.947, corresponding to Wright’s inbreeding coefficient (*F*) of 0.9 was simulated when considering partially selfing populations.

To assess BGS effects from unlinked selected sites, we simulated a 200 kb functional region experiencing purifying selection, and a 10 kb neutral region (no sites experiencing selection) from which mean *B*-values were observed across 100 replicates. A recombination rate of *r* = 0.5 was applied to the site at the breakpoint between the 200 kb selected region and the 10 kb neutral region so that the two regions are effectively unlinked, modelling the indirect effect of selected elements on other chromosomes. To test the prediction of *B* in the presence of pervasive interference (*B*′) implemented in the *Bvalcalc* ‘calculateB_hrì applied programming interface (API), we simulated a non-recombining region of 10^4^ sites experiencing purifying selection (see Text S4 for details).

### Bvalcalc code and figure generation

When making comparisons between results from *Bvalcalc* and simulated data, the simulation parameters precisely matched the corresponding *Bvalcalc* population genetic parameters utilized as input for the CLI and APIs. For example, the basic model parameters which we describe above were specified in *Bvalcalc* with the parameters input file available at: https://github.com/JohriLab/Bvalcalc/blob/main/tests/testparams/nogcBasicParams.py. The other tested parameters are available in https://github.com/JohriLab/Bvalcalc/blob/main/tests/testparams/. Note that a rescaling factor of 100 was applied, modifying the inputs to match the simulated values. Where the DFE consisted of a single selection coefficient, the ‘--constant_dfè flag was applied. Similarly, for incorporating demographic changes, the ‘--pop_changè flag was applied. The code used to generate figures and results, including the *B*-maps, is available at https://github.com/JohriLab/Bvalcalc/tree/main/manuscript.

### Comparison of B-maps to empirical data

#### D. melanogaster

The *D. melanogaster* r6.56 annotations were downloaded from FlyBase [49]. *PhastCons* elements [50] were downloaded from the UCSC genome browser [51]. See Daigle *et al.* (2026)[52] for details regarding annotation filtering. The *D. melanogaster* sex-averaged crossover rate map was obtained from Comeron *et al.* (2012)[47]. Population genetic parameters are available, along with relevant citations, in the *Bvalcalc* templates: ‘DroMel_Cds’, ‘DroMel_Utr’ and ‘DroMel_Phastcons’. Parameters include population size and mutation rate from Keightley *et al.* (2014)[46], mean crossover rate from Comeron *et al.* (2012)[47], gene conversion from Miller *et al.* (2016)[53], demographic history from Laurent *et al.* (2011)[54] and Kapopoulou *et al*. (2018)[55], the DFE of new mutations in exonic sites of single-exon genes from Johri *et al*. (2020)[14] was applied to coding sequences (CDS), and the DFE of untranslated region (UTR) and *phastCons* elements from Daigle *et al*. (2026)[52]. These gamma-distributed DFEs were split into nine non-overlapping uniform distributions logarithmically spaced using the ‘--gamma_dfè flag, corresponding to bins of homozygous selection coefficient (*<u>s</u>*): 0 to - 10⁻⁸, −10⁻⁸ to −10⁻⁷…-10⁻¹ to −1.

The empirical *D. melanogaster* SNP data was generated by Coughlan *et al.* (2022)[56]. The final VCF was filtered to only include newly sampled individuals from Mana Pools National Park in Zimbabwe that were classified as being in the “South 1” group in that work.

##### Humans

The GENCODE v38 comprehensive annotation for GRCh38, corresponding to Ensembl release 104 and incorporating HAVANA gene models, was filtered to retain CDS coordinates [51]. The human 100-way *phastCons* element coordinates [50] were downloaded from the UCSC genome browser. Recombination rates were obtained from the high-resolution, sex-averaged deCODE crossover map of Halldorsson *et al.* (2019)[57]. Spatial variation in non-crossover/gene conversion rate was represented using the map from Palsson *et al.* (2025)[58]. Population genetic parameters are available, along with relevant citations, in the *Bvalcalc* templates: ‘HomSap_Cds’ and ‘HomSap_Phastcons’. Parameters include population size and demographic history from Gutenkunst *et al.* (2009)[59], crossover rate from Kong *et al.* (2012)[59], mutation rate of 2.003861 × 10^−8^from Buffalo and Kern (2024)[35], gene conversion from Palsson *et al.* (2025)[58], CDS DFE from Huber *et al.* (2017)[61], *phastCons* DFE from Di *et al.* (2025)[62]. These gamma-distributed DFEs were split into nine non-overlapping uniform distributions logarithmically spaced using the ‘--gamma_dfè flag, corresponding to homozygous selection coefficient (*<u>s</u>*) bins: 0 to −10⁻⁸, −10⁻⁸ to −10⁻⁷…-10⁻¹ to −1.

Human variant data came from the 1000 Genomes Project [63], and we specifically used the GRCh38 integrated, phased biallelic SNP+INDEL call set generated in the GRCh38 reanalysis [64]; low-quality regions were excluded with the IGSR **“**strict**”** GRCh38 accessibility mask.

#### A. thaliana

The *A. thaliana* TAIR10 [65] annotation files were downloaded from Ensembl Plants [66] and filtered for CDS sites. The sex-averaged crossover map was obtained from Rowan *et al.* (2019)[67]. Population genetic parameters are available, along with relevant citations, in the *Bvalcalc* template: ‘AraTha_Cds’. Parameters include selfing rate from Platt *et al.* (2010)[68], population size and demography from Durvasula *et al.* (2017)[69], crossover rate from Rowan *et al.* (2019)[67], mutation rate from Weng *et al.* (2019)[70], gene conversion from Yang *et al*. (2012)[71], CDS DFE approximated from Gossmann et al. (2020)[72]. We converted the DFE into four non-overlapping uniform distributions corresponding to effectively neutral (0>2*N_anc_ s* >-1), weakly deleterious (−1>2*N_anc_ s* >-10), moderately deleterious (−10>2*N_anc_s* >- 100), and strongly deleterious ((−100>2*N_anc_ s*>-2*N_anc_*) mutations, present in proportions *f*_0_, *f*_1_, *f*_2_, *f*_3_ respectively. We assumed that 30% of new mutations in coding regions were synonymous (and belonged to the effectively neutral class), resulting in the following DFE for coding regions: *f*_0_=0.429, *f*_1_=0.065, *f*_2_=0.097, and *f*_3_=0.410.

The empirical *A. thaliana* variant data was downloaded from 1001 Genomes [73], filtered to only include individuals from the “south_sweden” group. Multiallelic and non-SNP sites were filtered out using *VCFtools* [74].

##### Quantifying the correlation of nucleotide diversity and *B*

We restricted analyses to putatively neutral sequences by (i) removing sites annotated as UTRs and conserved elements (*phastCons*) in the relevant reference genome sequence and (ii) treating coding sequence as neutral only at 4-fold degenerate positions. For *D. melanogaster*, regions containing additional functional annotations were also removed (as detailed in Daigle *et al*. 2026 [52]). We computed nucleotide diversity (the mean number of pairwise differences among all called chromosomes, divided by the number of callable sites) in non-overlapping windows of 1 Mb, 100 kb, 10 kb for each population using *scikit-allel* [75]. Windows with fewer than one neutral SNP were excluded. For each window, we took the mean *B* from the corresponding *B*-map and quantified the genome-wide correlation between *B* and nucleotide diversity using Pearson *R*^2^.

## Results and Discussion

### Comparison to simulated populations

To validate the effectiveness of our methods and implementation, we compared results from *Bvalcalc* to *B*-values observed from corresponding forward-in-time simulations (*SLiM* [45]). First, we tested the effect of background selection at neutral sites adjacent to a single 10 kb element experiencing purifying selection under a basic model with a DFE, constant population size, and no gene conversion or selfing (see Methods). We accurately recovered the mean effects of BGS observed in our simulations across 10,000 replicates (Figure 2a), using our implementation of Equations 2 and 3 in the *Bvalcalc* ‘genè module. Due to the assumptions involved in deriving Equation 3, *Bvalcalc* ignores the contribution of weakly deleterious mutations with 2*N_anc_sh* > −5 to BGS. We tested our predictions against simulations to ensure that this does not lead to substantial misinference when DFEs include a large proportion of mutations with selective effects below this threshold. We find that *Bvalcalc* effectively recovers the effects of BGS observed in simulations with a DFE consisting entirely of weakly deleterious (*f*_1_) mutations uniformly distributed between −1 > 2*N_anc_s* ≥ −10 (Figure S3).

**Figure 2.**
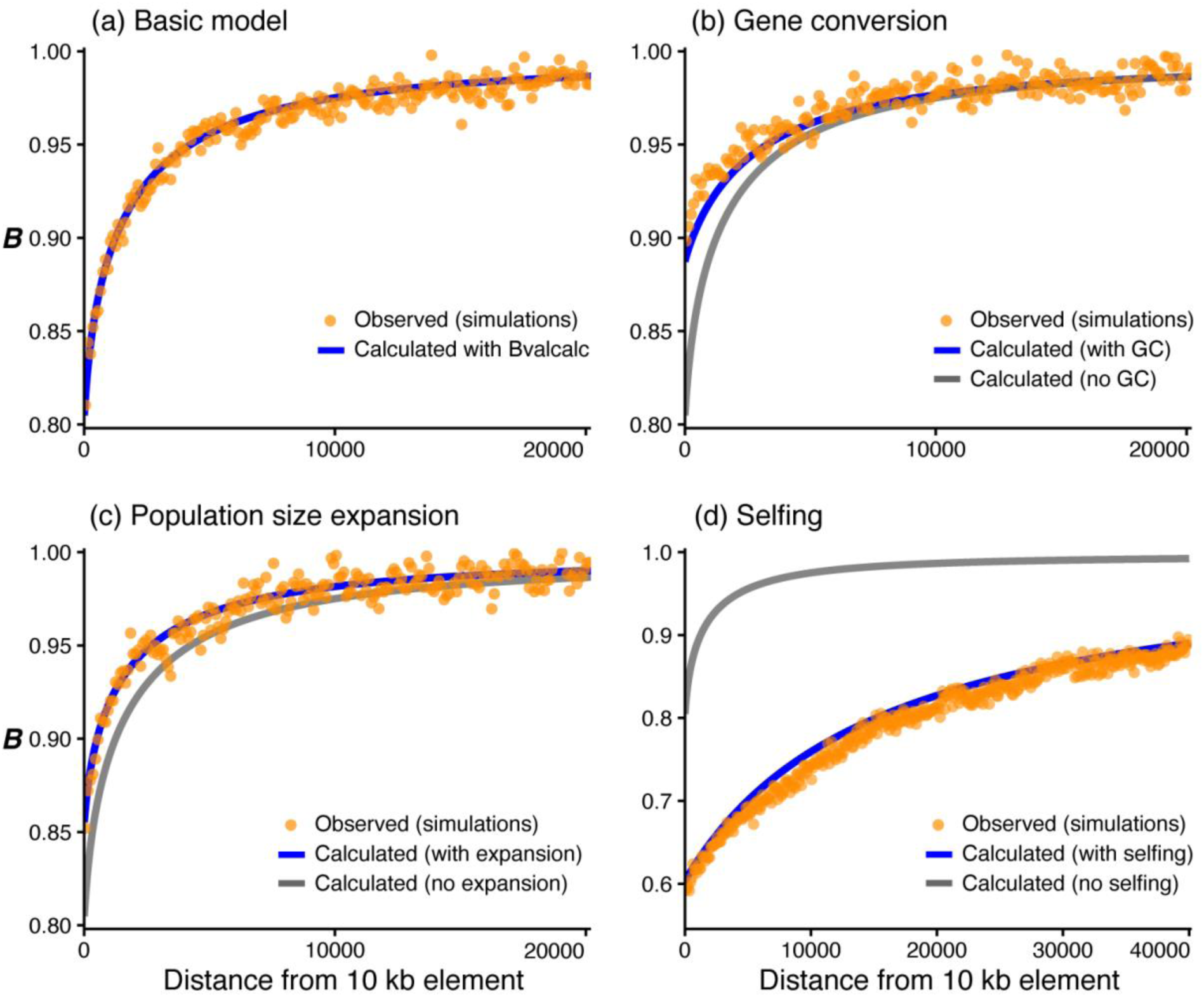
Recovery of nucleotide diversity at neutral sites (measured by *B*) with increasing distance from a 10 kb selected element experiencing purifying selection under the basic model (a), and with additional complexity (b-d). The orange dots represent observed results from simulations averaged over 10000 replicates. The lines represent predicted results using *Bvalcalc*’s ‘genè module: blue represents correctly specified parameters while grey represents the basic model in panel (a). Gene conversion (b) was modelled with 440 bp tract length and an initiation rate of *g* = 1 × 10^−8^per site/generation. The population size expansion (c) was a stepwise 5 × expansion 1*N_anc_* generations prior to sampling. The inbreeding coefficient (*F*) modelled for the selfing panel (d) was 0.9.

*Bvalcalc* offers several options for specifying a DFE, including a functionality to split a gamma distribution into 9 non-overlapping uniform distributions from neutral (*s* = 0) to homozygous lethal (*s* = −1). To ensure that this approach sufficiently approximates the gamma distribution, we compared simulations with mutations whose selective effects were drawn from a gamma distribution to the DFEs comprising uniform distributions. We found that splitting the gamma distribution into the 4 uniform distributions defined by *f*_0_, *f*_1_, *f*_2_, *f*_3_ led to an overestimation of *B*-values at nearby linked sites (Figure S4), underestimating the extent of BGS at sites close to the selected element. However, we found that splitting the DFE into 9 uniform distributions was generally effective at estimating the observed *B*-values (Figure S5); this is the default approach implemented in *Bvalcalc* for modelling a DFE when the ‘gamma_dfè flag is active. Note that users may specify an arbitrary number of non-overlapping bins to achieve greater DFE granularity as required with the ‘custom_dfè flag.

When gene conversion is added to the above model, the weakened effects of BGS are accounted for by *Bvalcalc* (Equation ST1.6; Equations ST2.1-2.6; Text S2) when accurately specified in the population parameters input file (Figure 2b). Note that *SLiM* models gene conversion tract lengths from a geometric distribution with a specified mean, while *Bvalcalc* models a fixed tract length. When a selfing rate of *F* = 0.9 (*S* = 0.947) was simulated, *Bvalcalc* effectively recovered the increased BGS effects (Figure 2d).

*B*-values following a 5-fold population size increase (*N_cur_* = 5 × *N_anc_*), 0.2*N_anc_*, 1*N_anc_* and 2*N_anc_* generations before sampling could also be accurately accounted for with Equation 5 (Figure 2c, Figure S6, Figure S7). However, we note that there remained deviations between the simulated results and expected values calculated with *Bvalcalc* after a 5-fold population size contraction (*N_cur_* = *N_anc_* / 5), 0.2*N_anc_* generations ago (Figure S8). We found somewhat greater deviations between simulated and calculated results at nearby sites when the contraction occurred 1*N_anc_* or 2*N_anc_* generations ago (Figure S9, Figure S10) as compared to when the change occurred more recently, 0.2*N_anc_* generations ago.

To further investigate this effect, we tested the 5-fold population contraction scenario 1*N_anc_* generations prior to sampling with different DFEs, which revealed that predicted *B*-values are more likely to deviate from simulated results when there are more weakly deleterious mutations present under a population size contraction (Figure S11, Figure S12). More specifically, our calculations overestimate the extent of BGS acting at sites close to functional elements. This is because after the population contraction, there is a decrease in the efficacy of selection (i.e., −2*N_cur_s* < −2*N_anc_s*), resulting in a higher proportion of effectively neutral and weakly deleterious mutations than present in the ancestral population. In fact, the extent of BGS generated from strongly deleterious mutations alone with selective strength 2*N_anc_s* = −100 (2*N_cur_s* = −20) was accurately estimated following the population contraction (Figure S13). These results (Figure S11, Figure S12, Figure S13) suggest that under a population contraction, the transition of mutations from 2*N_anc_s* < −5 to 2*N_cur_s* > −5 is poorly captured with Equation 5 as the effect of BGS by this class of mutations is not explained by Equations 2 and 3.

In all tested scenarios of population size contraction, estimates of *B*-values at neutral sites across the 1 kb region adjacent to the selected element are more accurately predicted with *Bvalcalc* if a constant population size equivalent to the ancestral size is assumed instead of decline (Table S1). In contrast, better results are obtained when specifying accurate demography in the *Bvalcalc* parameterization for all population expansion scenarios (Table S1). We therefore recommend assuming equilibrium in scenarios where the population may have undergone decline. Finally, the effect of a five-fold change in population size does not lead to exceptionally large deviations in *B*-values. For example, even in the simulated scenario that produced the least accurate predictions (a five-fold contraction 1*N_anc_* generations prior to sampling), the observed mean *B*-value (*B* = 0.84) in the 1 kb flanking region is comparable to the *Bvalcalc* estimates calculated assuming demography (*B* = 0.79) or assuming constant population size (*B* = 0.86).

We also validated the effects of BGS due to multiple selected elements at a site embedded in a small genomic region (200 kb including 10 equally spaced elements, each of length 10 kb) by comparing observations from simulations and corresponding *B*-maps obtained using *Bvalcalc’s* ‘region’/‘genomè modules (Figure 3). We find that under the basic model we can recover the observed diversity at intergenic sites using the equations implemented in *Bvalcalc* (Figure S14), confirming that *B*-values can be calculated precisely when assuming multiplicative effects from all conserved elements, where epistasis is absent and HRI is not pervasive. The addition of a variable recombination rate map leads to increased variability in *B*-values, which *Bvalcalc* estimated effectively (Figure 3).

**Figure 3.**
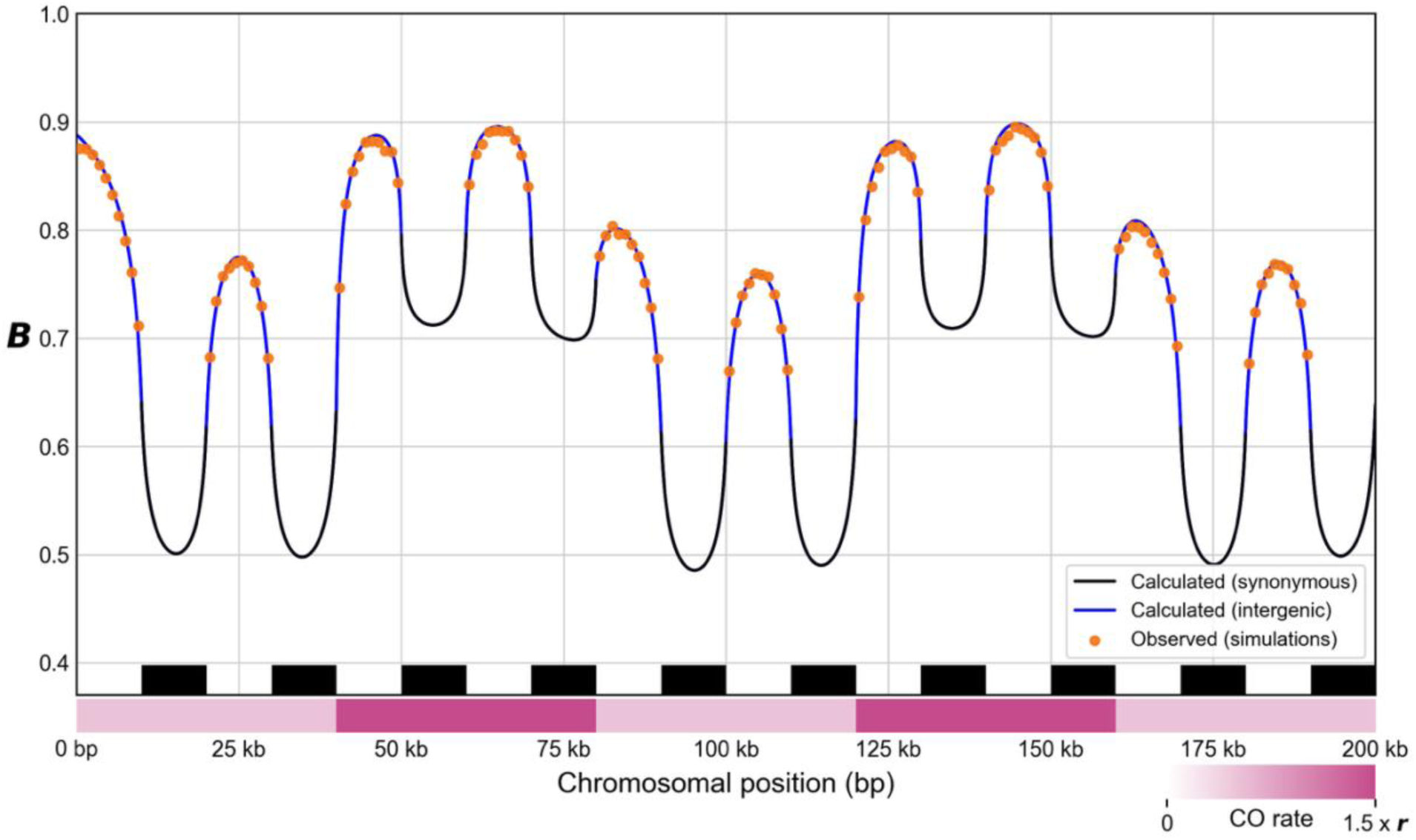
Predicted vs observed *B*-values across a 200 kb genome with 10 evenly distributed 10 kb selected elements (black segments at the bottom) experiencing purifying selection, including variation in crossover rate (pink-red segments at the bottom) across the region. The crossover rate varied relative to the mean rate (*r*) from *r* × 1.5 (dark pink) to *r* × 0.5 (light pink) in 40 kb intervals. The orange dots represent observed results from simulations averaged over 10,000 replicates. Lines represent predicted values obtained using *Bvalcalc*’s ‘region’ module: blue lines indicate *B*-values for neutral sites, while black lines indicate *B*-values for sites within selected elements.

We tested the effectiveness of Equation ST1.10 for modelling the unlinked effect of selected genomic elements on other chromosomes. We simulated the effects of BGS from 200 kb of unlinked selected sites experiencing purifying selection characterized by different DFEs on 10 kb of neutral sites. We compared the observed *B*-values on the 10 kb of neutral sites to corresponding calculations (Table 1). Equation ST1.10 overestimated BGS effects across all tested parameters, underestimating *B* by 12% for *s* = −0.1, *h* = 0.5, and by 76% for dominant lethal mutations (*s* = −1, *h* = 1), as it was derived assuming that *sh* ≫ −1 (see Appendix of [26]). We therefore extended the derivation relaxing this assumption, obtaining analytical expressions for *B* while accounting for unlinked effects of strongly deleterious mutations (Equation 4). Across all parameter combinations, the observed *B*-values from unlinked BGS were accurately recovered by Equation 4 for DFEs and for fixed selection coefficients (Table 1). Equation 4 is how *Bvalcalc*’s *B*-maps account for BGS from selected sites on other chromosomes; it is also available as an API.

**Table 1.** Comparison of predicted and observed (*i.e.*, simulated) effects of unlinked background selection generated by 200 kb of selected sites. Results averaged from 10 kb of neutral sites across 100 replicates are shown. Here, the simulated mutation rate was 1.2 × 10^−6^ per site per generation, with a population size of 10^4^. *f*_0_, *f*_1_, *f*_2_, and *f*_3_ represent the proportion of new mutations at selected sites that belong to the class of neutral, slightly deleterious, moderately deleterious, and strongly deleterious mutations respectively. Equation ST1.10 does not account for strongly deleterious mutations, while Equation 4 (implemented in *Bvalcalc*) does. For the final five rows, all new mutations at selected sites experienced a fixed selection coefficient.

| DFE ($f_1, f_2, f_3, f_4$) | Observed $B$ | Eq. ST1.10 | Eq. 4<br>(*Bvalcalc*) |
| --- | --- | --- | --- |
| Basic model (0.1, 0.2, 0.3, 0.4); $h=0.5$ | 0.89 | 0.82 | 0.89 |
| Strongly deleterious only (0, 0, 0, 1);<br>$h=0.5$ | 0.76 | 0.62 | 0.76 |
| $s = 0.1, h = 0.5$ | 0.92 | 0.91 | 0.92 |
| $s = 0.2, h = 0.5$ | 0.85 | 0.83 | 0.85 |
| $s = 0.5, h = 0.5$ | 0.74 | 0.62 | 0.74 |
| $s = -1, h = 0.5$ | 0.65 | 0.38 | 0.65 |
| $s = -1; h = 1$ | 0.62 | 0.15 | 0.62 |

Finally, we tested our diploid extension of the Becher and Charlesworth (2025)[36] model for non-recombining regions experiencing HRI effects using the ‘calculateB_hrì API under different selection parameters (see Text S4 for detailed description and discussion). In their model, the *B*-values accounting for HRI are referred to as *B*′. We accurately estimated *B*′ when simulating a DFE of a single constant selective effect (Table ST4.1) and when only a single class of deleterious mutations modelled by a uniform distribution was considered (*i.e.*, the class of weakly deleterious mutations only or moderately deleterious mutations only) using the coarse-graining method [33]. We underestimated *B*′ compared to simulations when using a DFE that included multiple classes of mutations (Table ST4.2), likely due to limitations in the ability of the coarse-graining approach to approximate a wider DFE with a single effective selection coefficient. Note that the effects of BGS generated by the class of strongly deleterious mutations did not result in much HRI and were thus always predicted well by our equations.

### *B*-maps in model organisms

As proof of concept, we used *Bvalcalc*’s ‘genomè module to construct *B*-maps across the autosomes of three model species using informed parameters from the literature: *H. sapiens* (HomSap) using coding sequences (CDSs) and *phastCons* elements [50], *D. melanogaster* (DroMel) using CDSs, UTRs and *phastCons* elements, and *A. thaliana* (AraTha) using CDSs only. Crossover rate maps and gene conversion initiation rate maps were used when available (see Methods). The *A. thaliana B*-map is, to our knowledge, the first published map of background selection for the species. For *A. thaliana* we only considered BGS effects from CDS sites; future work may improve on our map by accounting for non-coding conserved regions and by testing alternative parameters or inferring more accurate parameters.

*A. thaliana* exhibits the lowest mean *B*-value calculated using CDS sites only (*B̅* = 0.56 from CDS; Table 2), consistent with stronger effects of BGS under high selfing (*F* = 0.97; Platt et al. 2010 [67]). The *D. melanogaster* map also reveals strong effects of BGS at linked sites (*B̅* = 0.84 from CDS only; *B̅* = 0.64 from CDS + UTR + *phastCons*; Table 2), with pronounced within-chromosome variation in *B* driven by recombination rate heterogeneity (see Figure 4 for an example snapshot of a low-recombination region plotted by *Bvalcalc*). Humans have the highest *B̅* (0.87 from CDS only; 0.73 from CDS + *phastCons*), as expected. Notably, the contribution of BGS from unlinked sites to the mean *B*-value is greater for humans (41% of effects from unlinked sites; Table 2) than *A. thaliana* (11%) and *D. melanogaster* (22%), due to the substantially larger chromosome number.

**Figure 4.**
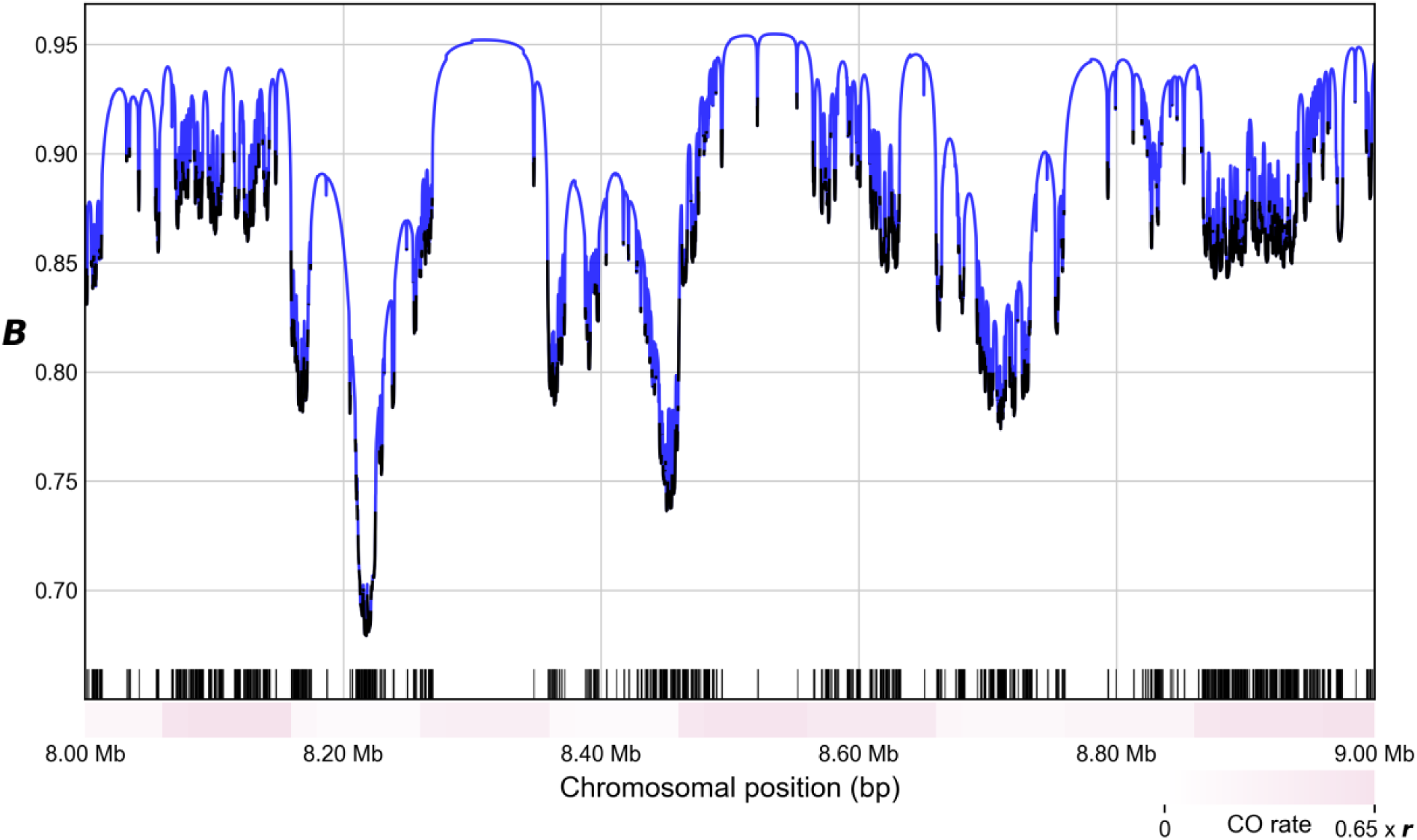
An example plot generated by *Bvalcalc* displaying a *B*-map across a 1 Mb region of interest on chromosome 2R of *D. melanogaster*. The *B*-map (only considering CDS) was generated using the ‘region’ module with the ‘--plot’ option. The blue lines indicate *B*-values for non-CDS sites, while black indicates *B*-values for sites within CDS elements. The black segments at the bottom represent selected elements (CDS in this case), and the pink-white segments represent variability in the local crossover rate. Note that the region shown here has a relatively low average rate of crossover to aid the visualization of changes in the *B*-map across the region.

**Table 2.** *B*-values averaged across autosomes (*B̅*) in three model species. “*B̅* (CDS)” denotes calculations using only coding sequences as targets of selection; additionally including untranslated regions (UTR*) and conserved intergenic elements (*phastCons*^), where modelled; *B̅_unlinked_* is the mean contribution from unlinked BGS (*i.e.,* effects from other chromosomes). *R*² values report genome-wide correlations between nonoverlapping windowed *B*-values and nucleotide diversity (*π*) at 1 Mb, 100 kb, and 10 kb scales. The total autosomal *B*-map length is the sum of the lengths of all autosomes over which the *B*-map was generated (and is slightly smaller than the sum of the actual lengths of all autosomal chromosomes).

| Species | $\bar{B}$<br>(CDS) | $\bar{B}$ (CDS +<br>UTR* +<br>*phastCons*^) | $\bar{B}_{unlinked}$ | Total $B$-<br>map<br>length | $R^2$<br>(1<br>Mb) | $R^2$<br>(100<br>kb) | $R^2$<br>(10<br>kb) |
| --- | --- | --- | --- | --- | --- | --- | --- |
| *Homo sapiens* | 0.87 | 0.72^ | 0.89^ | 2875 Mb | 0.48^ | 0.29^ | 0.14^ |
| *Drosophila melanogaster* | 0.78 | 0.61*^ | 0.92*^ | 108 Mb | 0.79*^ | 0.57*^ | 0.30*^ |
| *Arabidopsis thaliana* | 0.56 | - | 0.95 | 119 Mb | 0.44 | 0.26 | 0.15 |

\* $B$ -values were calculated including sites in UTRs.
^ $B$ -values were calculated including *phastCons* elements that were not already present in CDS or UTRs (when considered).

Genome-wide correlations between expected *B* and observed nucleotide diversity are strongest in *D. melanogaster* (e.g., *R*^2^ = 0.76 at 1 Mb scale; Table 2) and intermediate in *A. thaliana* (*R*^2^ = 0.43) and humans (*R*^2^ = 0.46 at 1 Mb). As window size shrinks from 1 Mb to 10 kb, correlations drop across all species, likely reflecting less precise local estimates of mutation rate, recombination rate, and genomic architecture, as well as greater variance in nucleotide diversity from fewer sites per bin (Table 2). We find that our *D. melanogaster B*-map exhibits similar high correlations to observed diversity as the map estimated by Elyashiv *et al.* (2016) [10] at 1 Mb scale (*R*^2^ = 0.76 for both maps, though note that Elyashiv *et al.* filtered out low-recombination regions). However, at finer scales, our *B*-map shows a higher correlation, *e.g.*, our map has *R*^2^ = 0.54 at 100 kb and *R*^2^ = 0.28 at 10 kb, while Elyashiv *et al.* report *R*^2^ = 0.42 at 100 kb scale and *R*^2^ = 0.23 at 10 kb. We attribute the accuracy of our *D. melanogaster B*-map to highly informed parameter choices, for instance, we assume the specific DFEs for the different categories of selected elements as inferred in Daigle *et al.* (2026)[52]. We note that values of *R*^2^ in humans were most sensitive to assumptions about recombination rate, with correlations between predicted and observed diversity being highest when fine-scale recombination rate was used (Figure S15).

To compare with previous methods, we directly compared the outputs of *Bvalcalc* human *B* maps with those generated by Buffalo and Kern (2024)[35]. Although that study evaluated many combinations of functional annotations and model settings, we selected their YRI *phastCons-priority* classic *B* map, in which *phastCons* elements were prioritized over coding regions and genes. This map provides the most direct comparison to *Bvalcalc* because both use classic BGS theory, whereas Buffalo and Kern’s primary *B*′ model is based on quantitative-genetic theory. We also included the more recent map from Barroso and Ragsdale (2026)[31]. This map was calculated using a mechanistic framework that predicts the effects of BGS using a two-locus model implemented in *bgshr*/*moments ++* [31], accounting for non-equilibrium demography. Notably, they also use pre-calculated parameters, including fine-scaled mutation and recombination rate maps, fine-scale DFEs for coding and noncoding regions under different levels of conservation to generate the *B*-map rather than jointly fitting the *B*-map and parameters. All maps were summarized in the same nonoverlapping 1 Mb windows. Across 2,456 common 1 Mb windows, the *Bvalcalc B*-map explained 47.5% of the variance in YRI diversity, compared with 44.8% for the Buffalo and Kern map and 54.1% for the Barroso and Ragsdale map (Figure S16). Note that the value of *R^2^* for the Buffalo and Kern map reported by us is much smaller than the value reported by them. Their reported value (63%) was obtained using code that calculates the Pearsons’ correlation coefficient (*R*), while we report the squared value, corresponding to *R^2^* (44.8%). Thus, *Bvalcalc* produces comparable outputs to preexisting methods in humans using independently specified inputs without the need for direct fitting of DFE or mutation rate parameters.

There are several possible factors that may explain why the correlation between the *B*-maps and empirical nucleotide diversity is not higher. First, the contribution of BGS from unlinked sites can be substantial, almost exceeding the effects of selection at linked sites (Table 2), leading to less relative variation across local windows. Second, the DFE across different annotation types and protein-coding genes is usually assumed to follow the same distribution. Differences in the DFE across genomic elements experiencing varying levels of evolutionary constraint could affect the local effects of BGS. Third, a fine-scale recombination map is essential to make accurate predictions and is unavailable for many model organisms, including *D. melanogaster* and *A. thaliana*. Lastly, temporal changes in rates of recombination can decrease correlation between predicted and observed diversity [76][77].

Our results indicate that the effects of background selection from linked sites are, on average, stronger than from unlinked sites across the genome (41% from unlinked sites; Table 2), which differs from the conclusions of Matheson and Masel (2024)[78]. They suggested that unlinked effects in humans would be stronger than linked effects. The differences in our conclusions may be due to different deleterious mutation rates, modelling a different DFE, genome architecture, or differences in the derivation of expressions for *B* from unlinked sites. We note that calculations with the equation used by Matheson and Masel (2024), which is equivalent to our Equation ST1.10 will be inaccurate when modelling under stronger selection coefficients (∼*sh* < −0.1), compared to Equation 4 as implemented in *Bvalcalc* (Table 1).

The *B*-maps we calculated represent a proof-of-concept and a starting point for more detailed analyses; improved *B*-maps for these species, and non-model species, may be generated with relative ease in the future using more carefully chosen or newly inferred population parameters with *Bvalcalc*. *B*-value maps across autosomes that we have calculated using template parameters are available for download at Zenodo: https://doi.org/10.5281/zenodo.17334918.

### Use-cases

There are wide-ranging potential applications of *Bvalcalc* for applied and theoretical population genetics. Constructing a *B*-map using the ‘genomè module is highly useful for developing an evolutionary baseline model for downstream analysis and inference [79]. For example, *SweepFinder2* [7] is a sweep detection method that can take a *B*-map as an input to account for the confounding effects of BGS on scans of loci that may have experienced recent positive selection (Figure 1; see our tutorial: https://johrilab.github.io/Bvalcalc/guides/sweepfinder2.html).

Demographic inference methods accounting for BGS are emerging, such as *PSMC*+ [80], which can use a *B*-map to correct for biases due to BGS in estimates of historical population size (Figure 1). Even for demographic inference methods assuming neutrality such as *dadi* [58] and *fastsimcoal2* [81], and coalescent methods such as *Relate* [82] and *MSMC* [83], a *B*-map is highly advantageous as one can filter for the most neutrally evolving sites (highest *B*-values) across the genome, which minimizes biases due to BGS (Figure 1; see our tutorial: https://johrilab.github.io/Bvalcalc/guides/demography.html).

Calculating *B*-values and *B*-maps with *Bvalcalc* can support and, in certain cases, circumvent the need for population genetic simulations, which can be computationally intensive. Simulating whole genomes is currently not feasible for most species without extensive rescaling, which can introduce biases to BGS effects (see our in-depth exploration of this issue in Marsh et al. 2026 [48]). Inferring *B*-maps with *Bvalcalc* offers a tractable alternative for modelling genome-wide effects of BGS.

*Bvalcalc* is a substantive advance over previous methods by integrating features into a flexible, user-friendly tool that can be applied to address a wide range of evolutionary questions. Other tools for *B-*value and *B-*map inference include: *calc_bkgd* which was the first widely used open-source software to calculate *B-*maps [27,37], *bgspy*/*Bprime* which accounts for HRI [35], *bgshr* which accounts for HRI and non-equilibrium demography [31], and *Binfer,* which accounts for non-equilibrium demography and partial selfing [24]. *Bvalcalc* uniquely accounts directly for the effects of gene conversion, which is becoming increasingly relevant with the release of new gene conversion rate maps [57,84] that can be utilized by *Bvalcalc*. Available tools currently provide minimal documentation and functionality at the level of *B-*value calculation. In the case of *calc_bkgd*, *bgspy*/*Bprime* and *Binfer*, *B*-value calculation is primarily an intermediate step toward fitting likelihood-based inference models to observed data. In contrast, *Bvalcalc* provides CLI modules with extensive features and documentation to help users quickly and effectively calculate BGS effects from known parameters, providing a comprehensive foundation for downstream analysis.

*Bvalcalc* offers rapid approaches for accurately estimating and comparing the BGS effects on nucleotide diversity caused by different population parameters. Using the ‘genè module, users can near-instantaneously test the effects of different parameters (mutation rate, recombination rate, gene conversion, DFE, selfing *etc.*) by visualizing the recovery curve of *B*-values with increasing distance from a single selected element. Users can efficiently visualize *B*-values in specific regions of the genome using the ‘region’ module (Figure 4) to better understand notable outliers and regions of particular interest. The ‘calculateB_unlinked’ API also enables users to estimate *B* from unlinked sites if the relevant population genetic parameters and the count of unlinked selected sites are known. We have included a rescaling parameter (set to 1 for empirical analysis) in the provided *Bvalcalc* parameters files which allows users to rapidly test the effects of rescaling on BGS to optimize their simulation strategies.

### Impacts of mis-specifying input parameters

As with any analytical model, misspecification of *Bvalcalc* input parameters will limit the accuracy of calculated *B*-values. This is particularly important to account for when applying *Bvalcalc* to non-model species for which accurate estimates of population parameters may not be available.

To provide users a basic understanding of the parameters’ sensitivity, we show results obtained from *Bvalcalc* under different crossover rates (Figure S17), selfing rates (Figure S18), and DFE parameters (Figure S19). Mis-specifying the mean crossover rate (*r*) or an incorrect crossover rate map will impact linked BGS effects. Under the basic model we find that reducing the crossover rate 2-fold from 0.5 × 10^−8^ to 0.25 × 10^−8^ reduced the calculated *B-*value at the position adjacent to the 10 kb selected region from 0.805 to 0.729, *i.e.,* by ∼10% (Figure S17). Selfing rate mis-specification has strong effects on BGS that persist over longer distances from selected sites compared to effects from mis-specifying the recombination rate (Figure S17, Figure S18), with an incorrect prediction of the mean diversity across the entire genome. For instance, with a Wright’s inbreeding coefficient that is mis-specified by ∼11% (*F*=0.8 vs 0.9), *B*-values are over or underestimated by ∼11% (i.e., *B*=0.84 vs 0.75 at 10 kb from the selected element in Figure S18). However, the slope of the recovery of diversity near selected elements is not substantially affected when the rate of selfing is mis-specified.

Mis-specifying the distribution of fitness effects defined by a gamma distribution has relatively smaller effects on the accuracy of *B*-value predictions at linked sites. Under the basic model with *Drosophila melanogaster*-like gamma distribution parameters [60], changing the mean selection coefficient impacted *B*-values at sites near the selected region (Figure S19). The impact of doubling or halving the mean selection coefficient decayed with increasing distance and was minimal at 20 kb from the selected region where *B* ∼ 0.971 under all three selection coefficients (Figure S19). In contrast, changing the shape parameter two-fold led to more persistent impacts on *B*-values with increasing distance: *B* = 0.971 (*shape* = 0.347), *B* = 0.980 (*shape* = 0.174), *B* = 0.962 (*shape* = 0.694) at 20 kb from the selected region (Figure S19).

Mis-specifying the demographic history of the population typically leads to minor deviations in *B*-values across the genome at a broader scale, consistent with findings from Mackintosh *et al.* (2026)[24]. However, at sites close to the selected element, levels of diversity are substantially influencedby nonequilibrium population history. Not accounting for population expansion may lead to an overestimation of the dip in diversity near functional elements caused by background selection. On the other hand, if the population has undergone a contraction, results assuming equilibrium reasonably approximate the dip in diversity near selected sites (Table S1 and Figures S8-S10). Both gene conversion and non-equilibrium demography affect levels of diversity near selected elements at smaller scales substantially more than at a broad genome-wide scale. Thus, previous methods that fit the recovery of diversity near functional elements to infer parameters of selection while assuming equilibrium and ignoring gene conversion may be biased.

A fold-change in the genome-wide mutation rate raises the *B*-value to the corresponding fold-change exponent (*i.e.*, doubling the mutation rate squares the *B*-value). Therefore, heterogeneity in the mutation rate across the genome would lead to overestimation of BGS effects caused by selected sites in low mutation rate regions and underestimation of BGS effects from sites in mutation rate hotspots.

We encourage users to consider how misspecification of input parameters will impact their results; this can be easily done using the ‘genè module with the ‘plot’ option active or using the ‘calculateB_unlinked’ API to check the effect of unlinked BGS by providing a count of unlinked sites under selection.

### Caveats and potential extensions

While we believe that *Bvalcalc* represents a step toward better inference of BGS effects, there remain caveats and substantial room for further extensions of the model. Hill-Robertson interference (HRI) between selected alleles increases diversity and thus the observed *B*-values (Table ST4.1, ST4.2; Text S4). HRI is not modelled by *Bvalcalc* by default. We expect that BGS effects will be overestimated (lower *B*) in regions experiencing HRI (usually regions of low recombination). To provide some functionality for users to account for HRI, we have implemented an API that performs calculation of *B*-values including HRI effects (*B*′) for non-recombining diploid genomes, using expressions from Becher and Charlesworth (2025)[36] extended with the coarse-graining method described by Good *et al.* (2014)[33] to model a DFE. Users should note that this *B*′ estimation method is an approximate method for modelling a DFE and is an incomplete solution for recombining genomes experiencing HRI (see Text S4 for full description and discussion). Given that accurate estimation of *B*′ accounting for HRI can rely on accurate *B* estimation prior to applying interference corrections [31,36], we hope that our extensions of the classic BGS model aid in the development of improved estimators of *B* accounting for HRI.

It’s important to highlight the effects that HRI may have on two common use-cases of a *B*-map calculated assuming no interference effects, *e.g.*, by *Bvalcalc*. Firstly, when using a *B*-map to select neutral sites for, *e.g.*, demographic inference, the regions with the lowest *B*-values (experiencing the strongest HRI effects) will be underestimated, though the relative ordering of *B*-values from lower-BGS sites (higher *B*) to higher-BGS sites (lower *B*) will remain largely unchanged. Thus, while the precise *B*-values may not be accurate for regions experiencing strong HRI, users may be confident that the higher *B*-value regions still typically contain the sites least affected by BGS across the genome, which are the best candidates for accurate demographic inference. Additionally, for selective sweep scans that use a *B*-map to develop a null-model, such as *SweepFinder2* [7], overestimating the effects of BGS would lead to an increased false negative rate as likelihood scores of true selective sweeps are reduced in low *B*-value regions when accounting for the effects of BGS (see Fig. 1 in DeGiorgio et al. 2016); however, the false positive rate is not expected to be affected. Nevertheless, downstream applications for which the precise *B*-value is essential (*e.g.*, when fitting the recovery of diversity to parameters of selection as done in Booker et al. 2021[11]) will be biased as *B*-values are underestimated by *Bvalcalc* when HRI is pervasive. For these cases, users should consider alternative software that may account for HRI effects and nonequilibrium demography, such as *bgshr* [31] and *Binfer* [24].

While *Bvalcalc* can model a single step-change in population size (Equation 5), it is not a comprehensive model of BGS under non-equilibrium demography. Equation 5, implemented in *Bvalcalc*, applies rescaling of a pre-computed *B*-value, assuming that the population has evolved under ancestral population-scaled parameters (*e.g., N_anc_* × *s*, *N_anc_* × *μ*, *N_anc_* × *r*, *N_anc_* × *g*), which shifted recently to a different population size [4]. However, this doesn’t consider the effect that non-equilibrium demography has had on the population-scaled parameters (*N_cur_* × *s*, *etc*.) which define the *B*-value itself, resulting in variability in the effect of demography on BGS over the genome [30], particularly in the presence of HRI. Under population size contraction, the transition of the proportion of new mutations from 2*N_anc_s* < −5, which is well captured by Equations 2 and 3 under our modelled scenarios, to 2*N_cur_s* > −5 leads to overestimation of BGS effects (Figure S11, Figure S12, Figure S13). Future work may extend Equation 5 to account for this overestimation in the absence of HRI, given a DFE and time since population size change. Further development of models and algorithms is needed to account for the interactions between BGS and complex demographic histories (see [31]). We view the current *Bvalcalc* implementation for modelling population size change as a useful starting point that we expect to be generally effective for many empirical systems that have experienced recent population size increases (*e.g.,* humans and *D. melanogaster*; Figure 2c), though it is less accurate under population size contractions (Figure S8, Figure S9, Figure S10) and is unlikely to hold under fluctuations involving many epochs or complex models including admixture.

There are several other caveats and potential extensions. *Bvalcalc* currently does not model variable mutation rates across the genome, though note that precise *de novo* mutation rate maps relevant for *B*-maps are difficult to obtain in many organisms [85,86]; nevertheless, accounting for mutation rate variation would be a useful extension. *Bvalcalc*’s ‘genomè and ‘region’ algorithms include some shortcuts to enable tractable estimation at single-base resolution; the discretising (chunking) of recombination rates and distances between the focal site and distant selected elements leads to some noise, though we expect this bias to be minor with minimal directionality (see Text S3, including Figure ST3.1, for details). Our extension of the classic BGS model to accurately account for gene conversion (Eq. ST1.6; Text S2) assumes a fixed tract length. However, *SLiM* models a geometric distribution of gene conversion tract lengths, and our results match relatively well when the mean is accurately specified (Figure 2b). We therefore expect *Bvalcalc*’s implementation to be generally effective under many tract length distributions, besides multi-modal distributions. *Bvalcalc* currently only supports diploid autosomes, though it is certainly possible to extend the underlying equations to account for sex chromosomes, haploids and polyploids.

### Conclusions

We have demonstrated the effectiveness of *Bvalcalc*’s prediction of nucleotide diversity with BGS, with functionality to estimate *B*-values for a wide variety of potential applications. We find that several factors, including gene conversion, selfing rates and BGS from unlinked sites, can all substantially impact local effects of BGS and *B*-values genome-wide, indicating that unified models of BGS are important for accurate inference. Further developments to BGS models are needed. However, we would like to stress the importance of approachability by developing accessible software, so that empirical biologists can account for BGS effects in a wide range of analyses involving non-model organisms. While we only provide *B*-maps inferred for model species as a proof of concept, *Bvalcalc* is a flexible tool that may be applied to non-model species for which *B*-maps are not yet available and where the genomic landscape of BGS is much less understood.

## Data Availability

*Bvalcalc* is available with documentation at https://johrilab.github.io/Bvalcalc/. To get started with *Bvalcalc*, see our tutorial which introduces the core command-line functionality: https://johrilab.github.io/Bvalcalc/guides/quickstart.html. *B*-value maps across autosomes constructed in this study using template parameters are available for download at Zenodo: https://doi.org/10.5281/zenodo.17334918. The code used to perform all analyses and generate results presented in this paper, including the *B*-maps, is available at https://github.com/JohriLab/Bvalcalc-manuscript.

## Acknowledgements

The research in this study was conducted using computational resources provided by ITS Research Computing at the University of North Carolina at Chapel Hill. The authors declare no conflicts of interest. We thank Brian Charlesworth for helping us implement the code provided in Becher and Charlesworth (2025). We thank members of the Johri lab, including Solomon Sloat and James Crescenzi, for testing *Bvalcalc* and providing feedback throughout its development. We thank Andy Kern and Vince Buffalo for confirming that the value of *R*^2^ reported in their manuscript corresponds to the Pearsons’ correlation coefficient (*R*).

## Funding

Research reported in this publication was supported by the National Institute of General Medical Sciences of the National Institutes of Health under award number R35GM154969 to PJ. The funders had no role in study design, data collection and analysis, decision to publish, or preparation of the manuscript.

## Supporting information

**Table S1.** Comparison of predicted and observed (*i.e.*, simulated) effects of background selection at linked neutral sites averaged across a 1 kb region adjacent to a 10 kb element experiencing direct purifying selection under the basic model with different simple demographic histories (single stepwise change) modelled in the simulations, averaged across 10000 replicates (*B_observed_*). Corresponding *Bvalcalc* results are shown assuming no population size change (*B_cur_* = *B_anc_*, ‘--pop_changè not active) and with accurately specified *Bvalcalc* population size change input (with ‘--pop_changè active). *B_cur_*/*B_anc_* indicates the fold-change from the ancestral size to current population size. The ancestral population size was *N_anc_* = 10^4^for all simulations. For example, the first row specifies the results obtained where a five-fold population size contraction occurred 0.2 × 10^4^ generations prior to sampling.

**Figure S1.** Recovery of nucleotide diversity at neutral sites (measured by *B* averaged in 100 bp bins across 10000 replicates) with increasing distance from a 10 kb element experiencing direct purifying selection under the basic model with a 13*N_anc_* generation burn-in (red), compared to a 14*N_anc_* generation burn-in (teal).

**Figure S2.** The difference between mean *B* averaged in 100 bp bins across 10000 replicates with increasing distance from a 10 kb element experiencing direct purifying selection under the basic model calculated from simulations with a 13*N_anc_* generation burn-in (red), compared to a 14*N_anc_* generation burn-in.

**Figure S3.** Recovery of nucleotide diversity at neutral sites (measured by *B*) with increasing distance from a 10 kb element experiencing direct purifying selection under the basic model with a DFE consisting only of weakly deleterious (*f*_1_) mutations uniformly distributed between −1 > 2*N_anc_s* ≥ −10.

**Figure S4.** Recovery of nucleotide diversity at neutral sites (measured by *B*) estimated with increasing distance from a 10 kb element experiencing direct purifying selection under the basic model with a gamma DFE matching *D. melanogaster* nonsynonymous CDS mutations inferred by Huber *et al.* (2017): 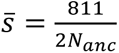, *shape* = 0.347. Here we simplified the gamma distribution into four bins by calculating the proportions of the gamma distribution within the classes defined by *f*_0_, *f*_1_, *f*_2_, and *f*_3_ which we used to parameterize our *Bvalcalc* calculations: *f*_0_ = 0.076, *f*_1_ = 0.093, *f*_2_ = 0.203, and *f*_3_ = 0.628. Note that the ‘--gamma_dfè flag was not used here. New mutations in the simulations sample from the gamma distribution directly (no simplification).

**Figure S5.** Recovery of nucleotide diversity at neutral sites (measured by *B*) estimated with increasing distance from a 10 kb element experiencing direct purifying selection under the basic model with a gamma DFE matching *D. melanogaster* nonsynonymous CDS mutations inferred by Huber *et al.* (2017): 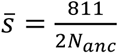, *shape* = 0.347. Note that *Bvalcalc* splits gamma DFE parameters into 9 non-overlapping uniform distributions (when ‘--gamma_dfè is active), while the simulations draw from the specified gamma distribution directly.

**Figure S6.** Recovery of nucleotide diversity at neutral sites (measured by *B*) with increasing distance from a 10 kb element experiencing direct purifying selection under the basic model with a stepwise 5 × expansion 0.2*N_anc_* generations prior to sampling.

**Figure S7.** Recovery of nucleotide diversity at neutral sites (measured by *B*) with increasing distance from a 10 kb element experiencing direct purifying selection under the basic model with a stepwise 5 × expansion 2*N_anc_* generations prior to sampling.

**Figure S8.** Recovery of nucleotide diversity at neutral sites (measured by *B*) with increasing distance from a 10 kb element experiencing direct purifying selection under the basic model with a stepwise 5 × contraction 0.2*N_anc_* generations prior to sampling.

**Figure S9.** Recovery of nucleotide diversity at neutral sites (measured by *B*) with increasing distance from a 10 kb element experiencing direct purifying selection under the basic model with a stepwise 5 × contraction 1*N_anc_* generations prior to sampling.

**Figure S10.** Recovery of nucleotide diversity at neutral sites (measured by *B*) with increasing distance from a 10 kb element experiencing direct purifying selection under the basic model with a stepwise 5 × contraction 2*N_anc_* generations prior to sampling.

**Figure S11.** Recovery of nucleotide diversity at neutral sites (measured by *B*) with increasing distance from a 10 kb conserved element experiencing direct purifying selection under the basic model with a stepwise 5 × contraction 1*N_anc_* generations prior to sampling; a DFE of only weakly deleterious mutations was modelled in the selected region: *f*_1_ only (a uniform distribution of −1 ≥ 2*N_anc_s* > −10).

**Figure S12.** Recovery of nucleotide diversity at neutral sites (measured by *B*) with increasing distance from a 10 kb element experiencing direct purifying selection under the basic model with a stepwise 5 × contraction 1*N_anc_* generations prior to sampling; a DFE of only moderately deleterious mutations was modelled in the selected region: *f*_2_ only (a uniform distribution of −10 ≥ 2*N_anc_s* > −100).

**Figure S13.** Recovery of nucleotide diversity at neutral sites (measured by *B*) with increasing distance from a 10 kb element experiencing direct purifying selection under the basic model with a stepwise 5 × contraction 1*N_anc_* generations prior to sampling; a DFE of only deleterious mutations of selective strength 2*N_anc_s* = −100 was modelled in the selected region.

**Figure S14.** Predicted vs observed *B*-values across a 200 kb genome with 10 evenly distributed 10 kb elements (black segments at the bottom) experiencing purifying selection, under the basic model with uniform recombination rate. The orange dots represent observed results from simulations averaged over 10,000 replicates. Lines represent predicted values obtained using *Bvalcalc*’s ‘region’ module: blue lines indicate *B*-values for neutral sites, while black lines indicate *B*-values for sites within selected elements.

**Figure S15.** Effects of successive parameter changes on *Bvalcalc* predictions for human chromosome 22. The left panel shows the squared Pearson correlation between predicted *B* and YRI nucleotide diversity across 27 common nonoverlapping 1 Mb windows. The right panel shows the corresponding *B*-maps along chromosome 22. We evaluated the effects of several parameter choices on model performance. The final model (orange) used the full-resolution deCODE sex-averaged recombination map, a mutation rate of 2.003861 × 10⁻⁸ per base pair per generation, a gamma distribution DFE split into 4 uniform distributions, and gene conversion. To assess the contribution of these choices, parameters were successively reverted from the final model: gene conversion was removed (green), the DFE was instead split into 9 bins as described in the Results. The mutation rate was reduced to 1.25 × 10⁻⁸ corresponding to base substitutions (blue); and the full-resolution recombination map was replaced by a 1-Mb average-rate map (grey). Population size change (*N*_anc_ = 7,300; *N*_cur_ = 14,474), CDS-priority annotation, and all other parameters were held constant.

**Figure S16.** Comparison of predicted *B*-values from *Bvalcalc* and two comparable methods with empirical nucleotide diversity in the Yoruba (YRI) population. Each point represents one of 2,456 nonoverlapping 1 Mb autosomal windows shared among all *B*-maps and passing a 25% callable-site coverage threshold. Black lines represent the fit of an ordinary least-squares regression to the data. We retained 2,456 autosomal windows that were represented in all four maps and had greater than 25% callable sites. For every map, we calculated the squared Pearson correlation between *B* and our estimates of YRI nucleotide diversity across this identical set of windows.

**Figure S17.** <u>The effect of mis-specifying the crossover rate (</u>*r*<u>) on</u> *<u>B</u>* calculated using *Bvalcalc* with increasing distance from a 10 kb element experiencing direct purifying selection under the basic model.

**Figure S18.** <u>The effect of mis-specifying the selfing rate (</u>*F*<u>) on</u> *<u>B</u>* calculated using *Bvalcalc* with increasing distance from a 10 kb element experiencing direct purifying selection under the basic model.

**Figure S19.** <u>The effect of mis-specifying a DFE provided by gamma distribution parameters on</u> *<u>B</u>* calculated using *Bvalcalc* with increasing distance from a 10 kb element experiencing direct purifying selection under the basic model. “Uncorrected” parameters refer to the basic model with a gamma DFE matching *D. melanogaster* nonsynonymous CDS mutations inferred by Huber *et al.* (2017): 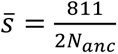, *shape* = 0.347. The mis-specified parameters scaled these values directly. For example, 2 × *shape* refers to *B*-values calculated with *shape* = 2 × 0.347 = 0.694, but otherwise the same parameters as “Uncorrected”.

## Supporting information captions

**Figure ST2.1.** Illustrative diagram describing the first subcase (i), with labels for the lengths of quantities relevant for calculating the probability of gene conversion events overlapping

(spanning) both the focal site and part of the element containing sites experiencing purifying selection (Equations ST2.1-ST2.6).

**Figure ST2.2.** Illustrative diagram describing the second subcase (ii), with labels for the lengths of quantities relevant for calculating the probability of gene conversion events overlapping (spanning) both the focal site and part of or the entire element containing sites experiencing purifying selection (Equations ST2.1-ST2.6).

**Figure ST3.1**. Illustrative diagram of the simplifications introduced by discretising recombination rates within chunks of uniform size (pink-red strip at the bottom) and combining distant selected elements (black blocks).

**Figure ST4.1**. Illustrative diagram of the difference between *B*′ within and around a non-recombining region experiencing strong HRI effects (green lines that account for HRI), and *B* assuming no HRI effects (black).

**Table ST4.1.** Background selection in a non-recombining region with constant selective effects.

**Table ST4.2.** Background selection in a non-recombining region with a DFE.

**Text S1.** Derivation of analytical expressions to obtain *B*-values

**Text S2.** Derivation of expressions accounting for complex gene conversion events.

**Text S3.** Description of *Bvalcalc*’s “genome” and “region” algorithms.

**Text S4.** Hill-Robertson interference effects and *B*′ calculation with *Bvalcalc* using the ‘calculateB_hrì API.

## Notes

https://github.com/JohriLab/Bvalcalc

https://github.com/JohriLab/Bvalcalc-manuscript

### Competing Interest Statement

The authors have declared no competing interest.

### Summary of Updates

Minor revisions of text addressing reviewer comments.

