## Supplementary material for "The *B*-value calculator: expected diversity with background selection": Text S1

### **Text S1**: Derivation of analytical expressions to obtain $B$-values

**Integration over a selected genomic element**

Here we obtain expressions for nucleotide diversity in the presence of background selection (BGS) relative to that under neutrality around a functional element assuming a DFE split into 4 bins of uniform distributions.

Let us assume that the functional element is of length $L$. The neutral site is $y$ distance from the end of the element and a selected site is $x$ distance (in the opposite direction) from the end of the element such that $z = x + y$ is the distance between the neutral and selected site. Let $B$ denote the nucleotide diversity in the presence of background selection (BGS) relative to that under neutrality. Then from equation 6 of Nordborg et al. (1996) we have:

$$B = \exp\left(-\iint E(t,z)\phi(t)\,dt\,dz\right) = \exp\left[-\int \phi(t)\left\{\int E(t,z)\,dz\right\}dt\right] \qquad \text{(ST1.1)}$$

Where $\phi(t)$ is the probability density function of $t$ $(sh)$ and $E$ is a function of the selection coefficient $(s)$, dominance coefficient $(h)$, rate of recombination, and gene conversion. Now we can evaluate the internal integral first (let's call it $F(t)$), which goes over the length of the genomic element. As we integrate over the functional element, $x$ goes from 0 to $L$ and $z$ goes from $y$ to $y + L$, where $y$ is a constant. That is,

$$F(t) = \int_y^{y+L} E(t,z)\,dz = \int_0^L E(t,x)\,dx$$

$$F(t) = \int_0^L \frac{1}{[t + (G[x+y] + R[x+y])(1-t)]^2}\,dx \qquad \text{(ST1.2)}$$

where $G \sim G(z)$ and $R \sim R(z)$ are the functions that describe the non-crossover (or gene conversion) and crossover rates between the neutral and selected sites respectively.

Assuming that $z$ is small and thus a linear recombination map is sufficient, we can make simplifying assumptions where $R(z) = rz$ and $G(z) = g$, a constant rate of initiation of gene conversion. In this case, the integral simplifies and we get:

$$G[x+y] + R[x+y] = g + r(x+y)$$

And

$$F(t) = \frac{Ut}{[t + (g + ry)(1-t)][t + (g + rL + ry)(1-t)]} \qquad \text{(ST1.3)}$$

This equation had previously been derived in Campos and Charlesworth (2019) and Johri et al. (2020).

**Accounting for multiple crossovers**:
As the value of $z$ increases, we can no longer assume that the rate of recombination linearly scales with $z$. But note that as $x$ (distance from the end of a selected element) will usually be small in realistic genomes where it likely represents an exon or a regulatory element, we can assume that crossover scales linearly with $x$ but not with $y$. For $y$, we can use Haldane's function (Haldane 1919), which does not account for crossover interference, but allows us to account for the effect of multiple crossovers. In this case,

$$R[z] = \frac{1 - e^{-2rz}}{2}$$

*i.e.*,

$$R[x + y] = rx + \frac{1 - e^{-2ry}}{2}$$

Substituting this in Equation ST1.2 and integrating, we now get:

$$F(t) = \frac{Ut}{[t + (g + C)(1 - t)][t + (g + rL + C)(1 - t)]} \qquad (ST1.4)$$

Where

$$C = \frac{1 - e^{-2ry}}{2}$$

**Accounting for gene conversion:**
Note that the effect of gene conversion is most important between closely linked sites and is almost negligible at long distances (Andalfatto and Nordborg 1998). Thus, in this case, it would be important to integrate over $x$ and account for the tract length ($k$).

From Frisse et al. (2001) and Langley et al. (2000), we have that the probability of the neutral and selected allele ending up in different haplotypes due to gene conversion can be captured by:

$$G[z] = 2gk\left[1 - e^{-z/k}\right]$$

Or

$$G[x + y] = 2gk\left[1 - e^{-(x+y)/k}\right]$$

Note that this expression is not easily integrable. Thus one possibility is to use the Taylor series approximation where $1 - e^{-x}$ can be approximated as *x* if *x* is small enough. This was also used in Andalfatto and Nordborg (1998), where they assume that the tract length *k* is fixed. In that case,

$$G[z] = \frac{gkz}{k} = gz \ when \ z << k$$

$$G[z] = gk \ otherwise$$

As these would easily allow for the expression to be integrable, substituting these in equation 4 and integrating with respect to $x$, we now get:
If $y + L << k$:

$$F(t) = \frac{Ut}{[t + (1 - t)(gy + C)][t + (1 - t)(g(y + L) + C + rL)]}$$

And otherwise:

$$F(t) = \frac{Ut}{[t + (gk + C)(1 - t)][t + (gk + rL + C)(1 - t)]} \qquad (ST1.5)$$

Now note that all equations ST1.3, ST1.4 and ST1.5 are of the following form:

$$F(t) = \frac{Ut}{[t + a(1 - t)][t + b(1 - t)]} \qquad (ST1.6)$$

Where $a = gy + C$ and $b = g(y + L) + rl + C$ when $y + L \;<< k$;
and $a = gk + C$ and $b = gk + rL + C$, otherwise.

Here $a$ and $b$ are constants and $a$ is generally not equal to $b$.

Although Equation ST1.6 will work well when $x$ is very small, other sub-scenarios need to be considered when a genomic element is involved. The constants $a$ and $b$ were therefore modified to account for other scenarios. For the purpose of continuity, these modifications are detailed in the supplement Text S2.

**Integration over a distribution of fitness effects**

In order to integrate Equation ST1.6 over the DFE, let us assume that $\phi(t)$ follows a uniform distribution between $t_1$ and $t_2$. Let us assume that

$$H = -\int \phi(t)\left\{\int E(t,z)\,dz\right\}dt$$

$$H = -\int_0^1 \phi(t)F(t)\,dt$$

$$H = -\int_{t0}^{t1} \phi_{0-1}(t)F(t)\,dt - \int_{t1}^{t2} \phi_{1-2}(t)F(t)\,dt - \int_{t2}^{t3} \phi_{2-3}(t)F(t)\,dt - \int_{t3}^{t4} \phi_{3-4}(t)F(t)\,dt$$

Where $t_i$ corresponds to $s_i h$ and $t_0 = 0$ and $t_4 = 1$. As $\phi(t)$ is uniform within each of these intervals, we can calculate

$$H_{ti,tj} = \int_{ti}^{tj} \frac{f_i F(t)}{(tj - ti)}\,dt$$

and

$$H = \frac{f_0}{t1 - t0}\int_{t0}^{t1} F(t)\,dt + \frac{f_1}{t2 - t1}\int_{t1}^{t2} F(t)\,dt + \frac{f_2}{t3 - t2}\int_{t2}^{t3} F(t)\,dt + \frac{f_3}{t4 - t3}\int_{t3}^{t4} F(t)\,dt$$

Using partial fractions, we can integrate the above and get the general form:

$$H_{ti,tj} = \frac{f_i}{(tj - ti)}\frac{Ua}{(1-a)(a-b)(tj-ti)}\ln\left(\frac{a + (1-a)tj}{a + (1-a)ti}\right) - \frac{Ub}{(1-b)(a-b)(tj-ti)}\ln\left(\frac{b + (1-b)tj}{b + (1-b)ti}\right) \qquad \text{(ST1.7)}$$

We can now calculate *B* by using equations ST1.1 and ST1.7 as $B = exp(-H)$.

**The case of no recombination**:

In the case of no recombination, $a = b = 0$ and $F(t) = U/t$. Thus,

$$H_{ti,tj} = \frac{f_i U}{(tj - ti)} ln\left(\frac{t_j}{t_i}\right)$$

**The case when *a*=*b*:**

Equation ST1.7 is valid when $a \neq b$. However, there will be scenarios when $a = b$. In that case, equation ST1.6 will be of the form:

$$F(t) = \frac{Ut}{[t + a(1-t)]^2}$$

Using partial fractions, we can write the above as:

$$F(t) = \frac{U}{(1-a)[a + (1-a)t]} - \frac{Ua}{(1-a)[a + (1-a)t]^2}$$

Now integrating over the distribution of fitness effects, as above, we get:

$$\int F(t)\,dt = \frac{U}{(1-a)^2}\left\{\ln|a + (1-a)t| + \frac{a}{a + (1-a)t}\right\}$$

and

$$H_{ti,tj} = \frac{f_i}{(t_j - t_i)}\frac{U}{(1-a)^2}\left\{\ln\left|\frac{a + (1-a)t_j}{a + (1-a)t_i}\right| + \frac{a}{a + (1-a)t_j} - \frac{a}{a + (1-a)t_i}\right\} \qquad \text{(ST1.8)}$$

**Accounting for simple changes in demographic history**:

**A step change in size**:
We assume that a population changes in size from $N_{anc}$ to $N_{cur}$ at time $T$ (which is in units of generations) in the past. Let us assume that $B_{anc}$ and $B_{cur}$ are the values of $B$ in the ancestral and current population respectively. Then using equation 1b from Johri et al. (2021), we can express $B_{cur}$ as:

$$B_{cur} = \frac{B_{anc}\left[1 + (R-1)e^{\frac{-T}{2N_{cur} \times B_{anc}}}\right]}{\left[1 + (R-1)e^{-\frac{T}{2N_{cur}}}\right]} \tag{ST1.9}$$

where,

$$R = \frac{N_{anc}}{N_{cur}}$$

Note that these derivations assume that $T$ is small enough that $B_{cur}$ is more or less constant across time 0 to $T$. Also note that we altered the time $T$ input variable from $2N_{cur}$ (in Johri et al. 2021) to simply take the number of generations (unscaled) as input, for a more intuitive user experience.

**Accounting for selfing in populations**:
To account for self-fertilization, we provide the `selfing` template with population genetic parameters modified by Wright's inbreeding coefficient (*F*). *F* can be calculated from the selfing rate (*S*) such that $F = S/(2-S)$ (Nordborg 2000). The template can be accessed from the CLI: ` Bvalcalc --generate_params selfing`. Our implementation includes a reduction of the population size to an effective selfing-adjusted population size, $N\_eff = N / (1 + F)$, a decrease in the crossover rate, $r\_eff = r \times (1 - F)$, a similar decrease in the gene conversion initiation rate, $g\_eff = g \times (1 - F)$ and a modified dominance coefficient, $h\_eff = h + F(1-h)$. These modified parameters are substituted in the equations used to calculate *B*.

**Calculating the effects of BGS from multiple linked functional elements**:
For $n$ different linked functional elements, the extent of BGS ($B_{\text{linked}}$) at a focal site will be given by $exp(-H1 - H2 - \cdots Hn) = exp(-H1) \times exp(-H2) \times \ldots \times exp(-Hn)$, i.e.,

$$B_{linked} = \prod_{i=1}^{n} B_i$$

where $B_i$ is calculated using the equations above.

**Background selection from unlinked sites:**

When sites are completely unlinked, $r = 0.5$, unlinked effects of background selection ($B_{unlinked}$) for diploids are given by:

$$B_{unlinked} \sim \exp(-8 \sum_i u_i t_i)$$

Where $u_i$ is the mutation rate to the deleterious variant at site $i$ and $t_i = s_i h_i$ is the heterozygous selection coefficient at each site and is assumed to be small. Note that here the contribution to unlinked BGS is being summed up over all unlinked selected sites in the genome.

Let us assume that there are $M$ chromosomes and $L_j$ is the length of chromosome $j$. Let us also assume for the sake of simplicity that $f_{sel}$ represents the proportion of selected sites on all chromosomes. Then, unlinked effects of BGS at chromosome $j$ would be

$$B_j \sim \exp\left(-8uf_{sel} \sum_{k=1, k \neq j}^{k=M} L_k \sum_i t_i\right) \quad \text{(ST1.10)}$$

Now assuming that $t_i$ follow a set of four nonoverlapping uniform distributions, $\sum_i t_i$ can be written as:

$$\int_{t=0}^{1} \varphi(t) t \, dt = \sum_{i=0}^{3} \frac{f_i}{t_{i+1} - t_i} \int_{t_i}^{t_{i+1}} t \, dt$$

Note that here $i$ represents each DFE class.

$$= \sum_{i=0}^{3} f_i \frac{t_{i+1} + t_i}{2}$$

Substituting this in Equation ST1.10, we have:

$$B_j \sim \exp\left(-8uf_{sel} \sum_{k=1, k \neq j}^{k=M} L_k \sum_{i=0}^{3} f_i \frac{t_{i+1} + t_i}{2}\right) \quad \text{(ST1.11)}$$

Or alternatively,

$$B_j \sim \exp\left(-8uf_{sel} \sum_{k=1, k \neq j}^{k=M} L_k \left\{\frac{f_0(t_1 + t_0)}{2} + \frac{f_1(t_2 + t_1)}{2} + \frac{f_2(t_3 + t_2)}{2} + \frac{f_3(t_4 + t_3)}{2}\right\}\right) \text{(ST1.12)}$$

Equation ST1.10 above assumes that $t_i$ is small. If we allow for $t_i$ to be larger, which is particularly relevant in organisms like Drosophila where the DFE is skewed towards strongly deleterious mutations, unlinked effects of BGS are given by

$$B_{unlinked} \sim exp\left[-\sum\nolimits_i \frac{8u_i t_i}{(1 + t_i)^2}\right] \quad \text{(ST1.13)}$$

The above was obtained using the Appendix in the paper by Charlesworth (2012). As before, unlinked effects of BGS at chromosome $j$ would be

$$B_j \sim \exp\left(-8uf_{sel} \sum_{k=1, k \neq j}^{k=M} L_k \sum_i \frac{t_i}{(1 + t_i)^2}\right) \quad \text{(ST1.14)}$$

Again, assuming that $t_i$ follow a set of four nonoverlapping uniform distributions, the summation over $t_i$ can be replaced by an integral:

$$\int_{t=0}^{1} \varphi(t) \frac{t}{(1+t)^2} dt = \sum_{i=0}^{3} \frac{f_i}{t_{i+1} - t_i} \int_{t_i}^{t_{i+1}} \frac{t}{(1+t)^2} dt \qquad \text{(ST1.15)}$$

Let us denote the integral by $I$. Then, we have

$$I = \int_{t_i}^{t_{i+1}} \frac{t}{(1+t)^2} dt$$

Using partial fractions, we get,

$$I = \int_{t_i}^{t_{i+1}} \frac{1}{(1+t)} dt + \int_{t_i}^{t_{i+1}} \frac{1}{(1+t)^2} dt$$

Integrating the above gives

$$I = \ln(\frac{1+t_{i+1}}{1+t_i}) + \left[\frac{1}{1+t_i} - \frac{1}{1+t_{i+1}}\right]$$

Substituting the above and Equation ST1.15 in Equation ST1.14, we get,

$$B_j \sim \exp\left(-8uf_{sel} \sum_{k=1,k\neq j}^{k=M} L_k \sum_{i=0}^{3} \frac{f_i}{t_{i+1} - t_i} \{\ln(\frac{1+t_{i+1}}{1+t_i}) + \left[\frac{1}{1+t_i} - \frac{1}{1+t_{i+1}}\right]\}\right) \qquad \text{(ST1.16)}$$
