## Supplementary material for "The *B*-value calculator: expected diversity with background selection": Text S2

### **Text S2**: Derivation of expressions accounting for complex gene conversion events

Integrating over a DFE and the length of a genomic element gives us equations of the form provided in Equation ST1.6 in the derivations Text S1, where $a = gy + C$ and $b = g(y + L) + rl + C$ when $y + L \ll k$; and $a = gk + C$ and $b = gk + rL + C$, otherwise. Here, $g$ is the rate of initiation of gene conversion, $k$ is a fixed tract length, $L$ is the length of the genomic element, and $y$ is the distance of the focal site from the end of the genomic element. Here we modify the terms "$a$" and "$b$" when $y < k$. Because both $a$ and $b$ have terms corresponding to crossover and gene conversion, let us denote the terms related to gene conversion by $G$. In particular, let

$$a = C + G_a[y]$$
$$b = C + rL + G_b[y]$$

The effect of gene conversion in breaking up linkage between a focal site on a chromosome, and sites under selection in a conserved element, can be described by several cases. The probability of gene conversion including any single site in the genome is $gk$ (ignoring edge cases within $k$ sites of chromosome ends). First, when $y \geq k$ any gene conversion event that includes a selected site cannot span the focal site and vice versa. This is because there are $k$ possible gene conversion tract initiation locations that include each site, and the maximum distance a gene conversion tract can span from *e.g.*, the focal site, is $k - 1$, assuming a fixed value of $k$ and no variability in $g$ within $k$ distance of the focal site. Thus, when the element and focal site are separated by $k$, the probability of gene conversion including the focal site to result in breaking linkage with a given site in the element, $G_a[z]$, is the same as the probability of gene conversion occurring for each site in the element to break linkage with the focal site $G_b[z]$. Thus $G_a[y] = G_b[y] = gk$ when $y \geq k$. This case is the same as that provided in the main derivations Text S1, see Equations ST1.6.

However, when $y < k$, a gene conversion tract may span both a selected site and the focal site for any selected sites within $k - y$ distance from the focal site, with linearly decreasing probability with increasing distance from the focal site (Figure ST2.1). When a gene conversion tract includes both the focal site and a selected site, linkage is maintained and so the "recombination" effect of gene conversion is nullified for the purposes of calculating $B$-values. Thus, when $y < k$, $G_a[y]$ is reduced from $gk$ by the mean proportion of the selected element that is also included in a gene conversion tract which maintains linkage, across the $k - y$ possible gene conversion tract locations that include the focal site and the selected element. Similarly, $G_b[y]$ is reduced from $gk$ by the probability that a gene conversion event which includes any selected site in the element also includes the focal site. Let $G_{null,a}[z]$ be the mean proportion of the element that is included in the $k$ possible gene conversion events that include the tract lengths, given that overlap (a tract including both a focal site and part of the element) is possible. Similarly, we can define $G_{null,b}[z]$ to be the probability that, given a gene conversion tract includes a selected site, it also includes the focal site. In that case, we can write:

$$G_a[y] = gy + g(k - y) \times (1 - G_{null,a}[y]) \tag{ST2.1}$$

$$G_b[y] = gk \times (1 - G_{null,b}[y]) \tag{ST2.2}$$

*Subcase (i)*: First, we will consider the case where the distance and length of the element are sufficient that a gene conversion tract could not span from the focal site, entirely across the element and include neutral sites on the opposite side of the element, *i.e.,* when $k \leq y + L$. In this case, the selected sites greater than $k$ base pairs from the focal site will never overlap and thus will not contribute to $G_{null,a}[y]$. However, overlap is possible for sites in the element that are $k - y$ bases from the focal site, *i.e.*, $(k - y)/L$ proportion of the total element. For the closest selected site at a distance $y$ bp from the focal site, the probability of overlap is $(k - y)/k$. For the most distant site that could be included in a gene conversion tract that also includes the focal site, the probability of overlap is $1/k$. As the probability of overlap decreases linearly for sites with increasing distance, we can obtain the mean probability of overlap for sites in the element that *can* overlap ($> k - y$ bp from the focal site), as the mean between the probability of overlap at the closest site ($y$ distance from focal site) and overlap at the furthest site in the element ($k$ distance from focal site). Thus, for the case depicted in Figure ST2.1, where $y \geq k$ and $k < y + L$:

$$G_{null,a}[y] = \frac{k-y}{L} \times \frac{\frac{k-y}{k} + \frac{1}{k}}{2} \textit{ when } k \leq y \textit{ and } k \leq y + L \qquad \text{(ST2.3)}$$

*Subcase (ii)*: When the element is close and short enough that it is possible for a gene conversion tract to span the focal site, the entire element, and neutral sites on the distant site, i.e., when $k > y + L$, the probability of overlap at the furthest site in the element is instead $(k - y - L)/k$ (see depiction in Figure ST2.2). Therefore,

$$G_{null,a}[y] = \frac{\frac{k-y}{k} + \frac{k-y-L}{k}}{2} \textit{ when } k \leq y \textit{ and } k > y + L \qquad \text{(ST2.4)}$$

We now get $G_{null,b}[y]$ for each subcase.

*Subcase (i)*: When $k \leq y + L$, the probability of overlap for sites further than $k$ distance from the focal site is zero, *i.e.*, for $L - k - y$ sites, while the proportion element that can overlap is $(k - y)\ /\ L$. The probability of overlap for the site at distance $k$ is 1, and the probability of overlap for the selected site closest to the focal site is $(k - y)/k$, and the mean probability of overlap for the $k - y$ sites in the element that can overlap with the focal site can be obtained by the mean of those probabilities. Thus, the probability across the entire element that when a gene conversion event includes part of the element, it also includes the focal site, is

$$G_{null,b}[y] = \frac{k-y}{L} \times \frac{\frac{k-y}{k} + \frac{1}{k}}{2} \textit{ when } k \leq y \textit{ and } k \leq y + L \qquad \text{(ST2.5)}$$

*Subcase (ii)*: When the element is close and short enough that it is possible for a gene conversion tract to span the focal site, the entire element, and neutral sites on the distant site, when $k > y + L$, the probability of overlap at the furthest site in the element is instead $(k - y - L)/k$.

$$G_{null,b}[y] = \frac{\frac{k-y}{k} + \frac{k-y-L}{k}}{2} when\ k \leq y\ and\ k > y + L \qquad \text{(ST2.6)}$$

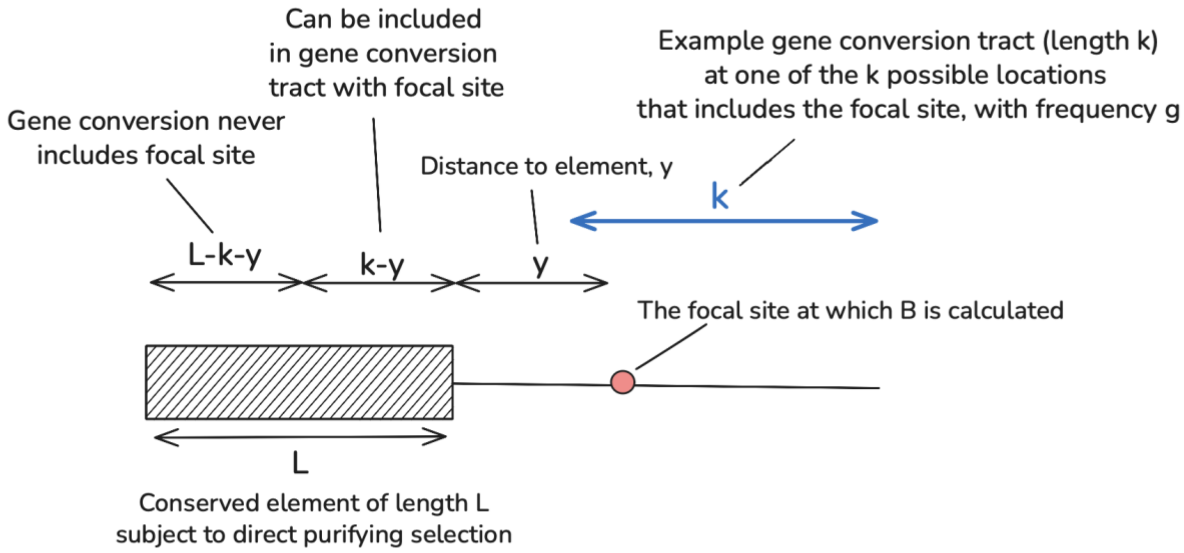


**Figure ST2.1. Illustrative diagram describing the first subcase (i), with labels for the lengths of quantities relevant for calculating the probability of gene conversion events overlapping (spanning) both the focal site and part of the conserved element containing sites experiencing purifying selection (Equations ST2.1-ST2.6).**

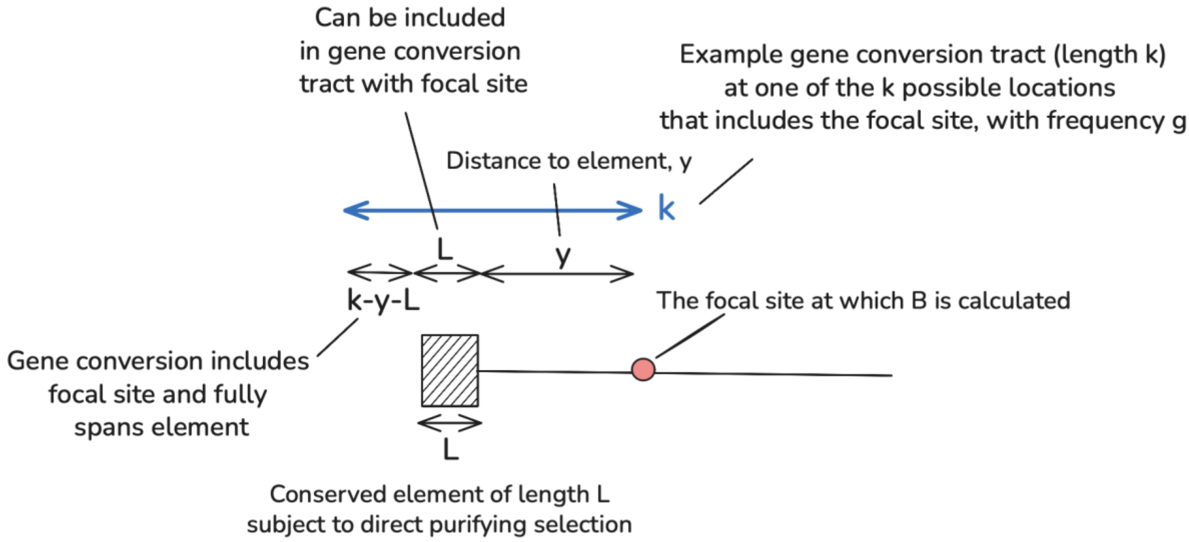


**Figure ST2.2. Illustrative diagram describing the second subcase (ii), with labels for the lengths of quantities relevant for calculating the probability of gene conversion events overlapping (spanning) both the focal site and part of or the entire conserved element containing sites experiencing purifying selection (Equations ST2.1-ST2.6).**
