## Supplementary material for "The *B*-value calculator: expected diversity with background selection": Text S3

### **Text S3**: *Bvalcalc* genome and region algorithms

*Bvalcalc*'s `genome` and `region` modules allow users to calculate *B*-maps at a single base pair resolution considering BGS effects from all conserved elements in the genome, linked and unlinked. The `genome` module calculates a *B*-map for all chromosomes in the provided BED/GFF annotation file. The `region` module calculates a *B*-map for only a specified region (in format CHR:START-END), while also considering genome-wide BGS effects on that region. The `region` module also allows for visualization of the *B*-map using `--plot` (*e.g.,* Figure 4). The `genome` and `region` modules share underlying code and so produce identical results for a given region, the only difference is the scope of the calculations.

Here, we describe some of the inner workings of *Bvalcalc*'s ability to tractably calculate a *B*-map, notably, breaking the genome into chunks for vectorized, multi-threaded computation. This is not an exhaustive description of the complete algorithm. For additional on-demand descriptions of the algorithm, consider providing the source code to a large language model capable of synthesizing the overall dependency graph.

See the relevant documentation at https://johrilab.github.io/Bvalcalc/modules/region.html, https://johrilab.github.io/Bvalcalc/modules/genome.html.

#### Calculating *B* for each site in a chunk of the genome

For each site in the genome, the contributing effect on *B* is calculated relative to every conserved element in the genome using the `calculateB_linear` (default) or `calculateB_recmap` (if a variable recombination rate map is provided) APIs. The contributing effects of BGS from all conserved elements are combined multiplicatively to obtain the *B*-value at a focal site (see the derivations Text S1). Thus, *B*-values are computed essentially from scratch for each site in the genome, albeit with some shortcuts.

There are a few key simplifications that avoid large increases to computation time with increasing number of conserved genomic elements. First, BGS from unlinked sites is calculated only once per chromosome using the total count of conserved sites in all other chromosomes, it is applied to all sites in the focal chromosome uniformly. Secondly, to enable vectorization of *B*-value calculations using *NumPy* arrays, the genome is split into contiguous chunks of uniform size (max 20 kb by default), and each chunk is processed independently using multi-threading. By default, the chunk size is dynamically chosen by an algorithm by the total number of conserved elements when a chromosome is loaded to avoid excess memory usage, though users may override this and set it manually with `--chunk_size`.

The recombination rates (gene conversion and crossover), when provided, are simplified so that only the mean rates per chunk are considered. Thus, if half of a chunk is specified as having the mean crossover rate (*r*), and the other half is specified as having a halved recombination rate relative to the mean (`0.5 * r`), *Bvalcalc* will simplify the entire chunk to have a uniform recombination rate of the mean rate, `0.75 * r` (Figure ST3.1). This is not biologically accurate, though vastly improves tractability and is expected to not lead to substantial biases for most analyses for which the chunk size will be much smaller than the intervals between recombination rate changes.

Next, the *B*-values for a given focal chunk are processed. The first step is to partition the remaining chunks in the chromosome into two categories, the "precise" chunks which are chunks directly adjacent to the focal chunk (3 chunks on either side of the focal chunk by default, which can be changed with `--precise_chunks`), and the "distant" chunks (greater than 3 chunks away from the focal chunk). *B*-values are calculated using the precise distance from and lengths of each conserved element in the "precise" chunks region. This is to ensure that the closest elements, which may have the strongest local effect of BGS, especially from moderate and weakly deleterious mutations, are accurately calculated. However, it is not practical to calculate *B* efficiently across large regions, for *e.g.*, 100 Mb of sites in a human chromosome, which may have ~500,000 conserved phastCons elements. To avoid this, the conserved elements in "distant" chunks (all non-"precise" chunks on the same chromosome) are simplified, so that *B* is calculated only once per distant chunk (Figure ST3.1). This simplification is done by only calculating *B* once using the length of the sum of all conserved sites within a given distant chunk with a distance calculated as though it's in the centre of the chunk. Thus, many conserved elements scattered across a 20 kb chunk that is greater than 60 kb (precise chunks, 3 * 20 kb chunk size) are simplified to represent a single conserved element placed directly in the centre of that chunk (Figure ST3.1). Note that each focal chunk will have a different set of "precise" chunks and "distant" chunks corresponding to its relative position.

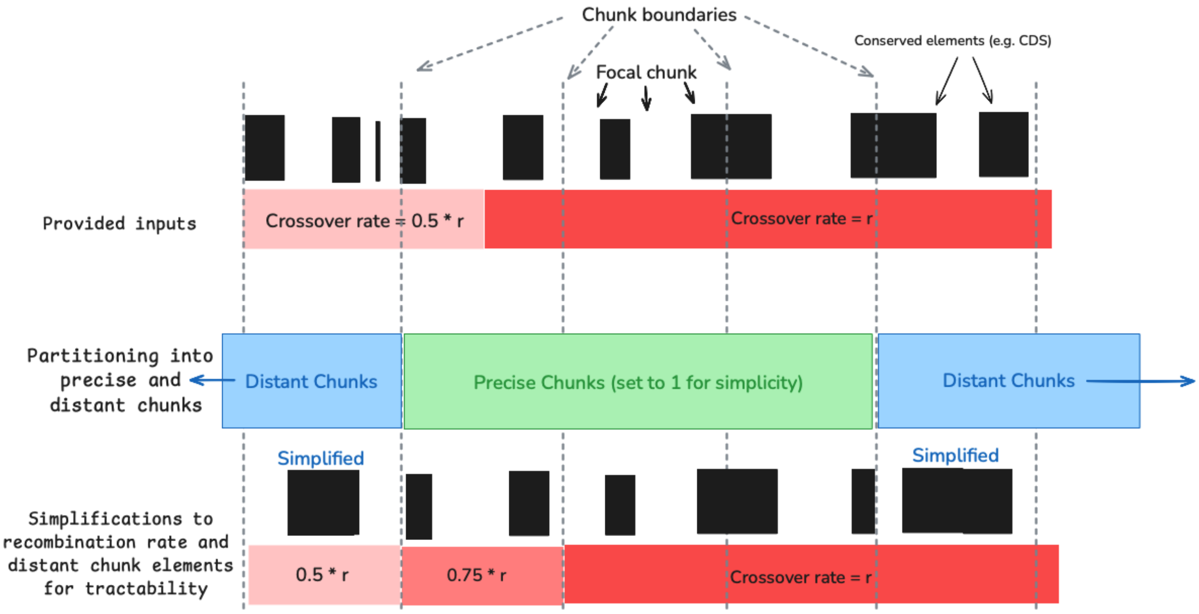


**Figure ST3.1. Illustrative diagram of the simplifications introduced by discretising recombination rates within chunks of uniform size (pink-red strip at the bottom) and combining distant conserved elements (black blocks).** Here, `--precise_chunks` is set to 1, indicating that only the focal chunk and the adjacent chunks are kept for precise calculations of length and distance to conserved elements, while chunks two or more chunk lengths away will be treated as distant chunks, with all conserved elements combined into a single central element. Note that by default, *Bvalcalc* has `--precise_chunks`

set to 3, which means that the three flanking chunks on each side are not simplified (7 chunks total including the focal chunk).

The only drawback of this simplification approach of distant chunks is that it does not calculate the distance to each conserved element precisely (Figure ST3.1). We justify this by assuming that minor deviations of distance to conserved elements are not important for distant sites, especially when the biases are introduced relatively evenly (some are shifted closer, others further). Nevertheless, this is the cause of the small stepwise changes in *B*-values between chunks that is observable in plots of *B*-maps generated by *Bvalcalc*. We however consider it a necessary drawback of the approach to enable efficient analysis of whole genomes. If this is a concern for your analysis, consider widening the "--precise_chunks" window. Users may choose to set it to cover all chunks on a chromosome to avoid this simplification and perform precise calculations for all elements, which may be computationally feasible for small chromosomes with few genomic elements.

#### Updating the overall *B*-map

Once the *B*-values for each site in each chunk are calculated from elements in all "precise" and "distant" chunks, a master array of the entire chromosome is updated. Processing of chunks continues by multi-threading until *B*-values for all chunks have been updated in the master array. When users export a *B*-map, the output bin size specified will be used to obtain the mean *B*-value for each bin across the region. For example, `--out_binsize 1000` outputs a *B*-map across the entire region with mean values for each non-overlapping bin of size 1000 bp. Users may set `--out_binsize 1` to retrieve the precise per-base resolution, though note that this will often produce an enormous output file size.
