## Supplementary material for "The *B*-value calculator: expected diversity with background selection": Text S4

### **Text S4**: Hill-Robertson interference effects

*Bvalcalc* calculates $B$ assuming independent dynamics of selection across selected loci. In other words, the interactions between selected alleles are not modelled. Hill-Robertson interference effects (HRI; Hill and Robertson 1966; Felsenstein 1974) occur when sufficient selected mutations are segregating concurrently, particularly while in linkage disequilibrium. HRI acts by increasing genetic diversity (relative to that under background selection only) and decreasing the efficacy of selection at interfering loci, which can change fixation probabilities of selected alleles and the rate at which deleterious mutations are purged from the population, particularly in gene-dense, low-recombination genomic regions. Specifically, HRI reduces the relative decrease in diversity due to purifying selection at linked sites as compared to background selection (BGS) alone when calculated using the Nordborg (1996) model ($B$), as is calculated by the primary modules of *Bvalcalc*. Relative diversity under background selection, adjusted for interference effects, is denoted as $B'$. Calculating precise estimates of $B'$ across the genome, while obviously valuable, is outside the current scope of *Bvalcalc*, which focuses on accurate and efficient estimation of $B$. Nevertheless, we have implemented an API in *Bvalcalc* to calculate $B'$ for a non-recombining region experiencing HRI from the model developed in Becher and Charlesworth (2025) which provides some tooling to approximate $B'$ in specific regions experiencing HRI, albeit with major caveats.

#### API implementation of $B'$ calculation

The `calculateB_hri` function available in the API (hereafter referred to as the API) requires users to input the cumulative length of selected sites in the focal region, $B$ from sites external to the non-recombining region (including from other chromosomes), and a *Bvalcalc* params file with population genetic parameters describing the population of interest. The haploid model is described in Becher and Charlesworth (2025), our implementation includes two major changes. First, we extended the model to diploids with changes including doubling the region-wide deleterious mutation rate ($U$), doubling and scaling selection by the dominance coefficient ($t = sh$). Second, we extended the model to a DFE using the coarse-graining method proposed by Good et al. (2014) which we apply to weakly selected ($-1 < 2N_es \leq -10$) and moderately selected ($-10 < 2N_es \leq -100$) mutations only to estimate a single selective strength to use for the calculations; we assume that nearly neutral mutations (present in $f_0$ proportion) are neutral and strongly deleterious mutations ($-100 < 2N_es$) mutations are too rapidly purged to contribute to interference effects (see * in Table ST4.2). Here, we will describe only the modifications of Becher and Charlesworth (2025) for convenience, though see Becher and Charlesworth (2025) for details regarding the derivation.

**Calculating the effective selection coefficient:**

Becher and Charlesworth's derivation of predicted $B'$ (i.e., effects of background selection accounting for interference) assumes a single selection coefficient. In order to implement a DFE of deleterious mutations, we used the method suggested by Good et al. (2014) to calculate an effective selection coefficient, or rather $t_{eff}$ ($hs_{eff}$) where $s$ represents the fitness disadvantage of the mutant compared to the wildtype. We assume that the DFE can be discretized into 4 non-overlapping uniform distributions comprising effectively neutral ($0 \leq 2Ns < 1$), weakly deleterious ($1 \leq 2Ns < 10$), moderately deleterious ($10 \leq 2Ns < 100$), and strongly deleterious ($100 \leq 2Ns$) mutations present in proportions $f_0$, $f_1$, $f_2$, and $f_3$ respectively. Let $\gamma_i(= 2Ns_i)$ represent the boundaries of the discretized distributions, i.e., $\gamma_1 = 1$, $\gamma_2 = 10$ and $\gamma_3 = 100$. In that case, $t_{eff}$ is defined as

$$t_{eff} = \sqrt{\frac{\frac{f1}{f1+f2} \times h^2 \times E_1}{4N_0^2} + \frac{\frac{f2}{(f1+f2)h}^2 \times E_2}{4N_0^2}} \quad \text{(ST4.1)}$$

where $N_0$ is the ancestral population size scaled by the $B$-value generated due to direct purifying selection at sites *not* within the non-recombining region, calculated for any site within the focal HRI region. That is, $N_0 = B_{distant} \times N_{anc}$. Thus, within the focal HRI region, $B_{distant}$ reflects the $B$-value calculated using equations detailed in the derivations Text S1 and the Methods, from all selected elements genome-wide, excluding the conserved sites within the HRI region. Additional terms are defined as

$$E_1 = \frac{({\gamma_1}^2 + \gamma_1 \times \gamma_2 + {\gamma_2}^2)}{3} \quad \text{(ST4.2)}$$

$$E_2 = \frac{({\gamma_2}^2 + \gamma_2 \times \gamma_3 + {\gamma_3}^2)}{3} \quad \text{(ST4.3)}$$

**Obtaining expected diversity under interference**:

As defined by Becher and Charlesworth, $\gamma = 2N_0 \times t_{eff}$, $U = 2\mu L_{del}$, $\alpha_2 = 2N_0U$, $k = 1$, where μ is the *de novo* mutation rate, $L_{del}$ is the sum of selected sites in the HRI region, and $k$ represents the mutation bias from one allele to another (which we assume to be 1). Note that these parameters are defined slightly differently from Becher and Charlesworth in order to account for diploidy. A modified version of their Equation 7 was used to obtain expected $B$ (which we call $B_{coal}$ here) by numerically solving the following equation by root finding (grid bracketing over $B \in (0,1]$ followed by linear interpolation / a single secant step):

$$-\ln(B_{coal}) = \frac{U \times (1 - e^{-\gamma B_{coal}})^3}{2t \times (1 + ke^{-\gamma B_{coal}})^3} \quad \text{(ST4.4)}$$

With this solution in hand, $B'$ can be calculated for sites in the non-recombining region under HRI by their Equation 12, which is reproduced here as

$$B' = B_{coal} \int_0^\infty \exp\left(-B_{coal} I(x)\right) dx \quad \text{(ST4.5)}$$

where

$$I(x) = \int_0^x g(s) ds \quad \text{(ST4.6)}$$

where

$$g(x) = \exp(c[1 - e^{-dx}]^2) \tag{ST4.7}$$

where

$$c = \frac{\alpha_2}{2\gamma} \times \left[\frac{1 - e^{-\gamma B_{coal}}}{1 + ke^{-\gamma B_{coal}}}\right]^3 \tag{ST4.8}$$

and

$$d = 2\gamma B_{coal} \times \frac{1 + ke^{-\gamma B_{coal}}}{1 - e^{-\gamma B_{coal}}} \tag{ST4.9}$$

In our implementation we truncate the upper limit of the integral in Equation ST4.5 from infinity to 100 as it saturates quickly, and the integral is approximated numerically by the trapezoid rule.

#### Comparison to simulated results

Using the API implementation of Equation ST4.5, we compare our calculated results to simulations of diploid non-recombining regions ($r = 0, g = 0$) with varying parameters of selection. Table ST4.1 indicates we can effectively predict observed BGS effects in diploid populations experiencing HRI when no recombination is present and selective effects are constant. In essence, Table ST4.1 confirms the results from Becher and Charlesworth (2025) and shows our extension to diploids is effective.

**Table ST4.1. Background selection in a non-recombining region with constant selective effects.** Observed $B$ is the mean diversity compared to neutral simulations collated from 100 replicates. Expected $B'$ was calculated using the `calculateB_hri` function in the *Bvalcalc* API. Expected $B$ was calculated using the `calculateB_linear` function in the *Bvalcalc* API and represents the expected effects of background selection in the absence of interference effects. $L_{del} = 10^4$ selected sites, $\mu = 3 \times 10^{-7}$, $h = 0.5$, $N_0 = 10^4$.

| $2Ns$ | Observed $\pi/\pi_0$ | Expected $B'$ (accounting for interference; API) | Expected $B$ (assuming no HRI) |
|---|---|---|---|
| 1 | 0.70 | 0.71 | 0.00 |
| 3 | 0.48 | 0.48 | 0.00 |
| 5 | 0.40 | 0.39 | 0.00 |
| 10 | 0.31 | 0.29 | 0.00 |
| 20 | 0.24 | 0.21 | 0.00 |
| 40 | 0.19 | 0.17 | 0.05 |
| 80 | 0.24 | 0.24 | 0.22 |
| 160 | 0.46 | 0.47 | 0.47 |
| 320 | 0.66 | 0.69 | 0.69 |

Extending the model further to include the coarse-graining method described by Good et al (2014) to accommodate a discretised DFE produced mixed results (Table ST4.2). We find that when we only simulate a single mutation class ($f_1$ only or $f_2$ only), $B'$ recovers observed $B$ well. In addition, we show that the $f_3$ class can be ignored for $B'$ calculations, as inputting additional $f_3$ mutations does not change the observed or expected $B$ and $B'$ (see * in Table ST4.2).

**Table ST4.2. Background selection in a non-recombining region with a DFE.** The DFE column represents the proportions of different classes of deleterious mutations. Observed $B$ is the mean diversity compared to neutral simulations collated from 100 replicates. Expected $B'$ was calculated using the `calculateB_hri` function in the *Bvalcalc* API. Expected $B$ was calculated using the `calculateB_linear` function in the *Bvalcalc* API and represents the expected effects of background selection in the absence of interference effects. $L_{del} = 10^4$ selected sites, $\mu = 3 \times 10^{-7}$, $h = 0.5$, $N_0 = 10^4$.

| $DFE\ (f0, f1, f2, f3)$ | Observed $\pi/\pi_0$ | Expected $B'$ (accounting for interference; API) | Expected $B$ (assuming no HRI) |
|---|---|---|---|
| 0, 1, 0, 0 | 0.38 | 0.36 | 0.00 |
| 0, 0, 1, 0 | 0.20 | 0.18 | 0.05 |
| 0, 0, 0, 1 | 0.97 | N/A^ | 0.97 |
| 0.1, 0.2, 0.3, 0.4 | 0.37 | 0.31 | 0.06 |
| 0.1, 0.2, 0.3, 1.4* | 0.37 | 0.31 | 0.06 |
| 0, 0.8, 0.2, 0 | 0.33 | 0.19 | 0.05 |
| 0, 0.2, 0.8, 0 | 0.23 | 0.17 | 0.01 |

*This DFE sums to 2 instead of 1 to reflect the increased mutation rate. In essence this is the same DFE as above, but with an additional $3 \times 10^{-7}$ mutations from the $f_3$ class. Here we show that changing the $f_3$ proportion does not lead to substantial differences in $B$ or $B'$, expected or observed, indicating we can ignore the $f_3$ class when calculating $B'$.
^As $B'$ is only calculated using the $f_1$and $f_2$ class mutations, $B'$ cannot be calculated here.

For DFEs combining different mutation classes, observed $B$ deviates from $B'$, e.g. the calculated $B'$ underestimates diversity by 42% compared to simulated results when the DFE is 80% $f_1$ and 20% $f_2$ (Table ST4.2). This is a shortcoming of the coarse-graining method using the root mean squared of selection coefficients, which is only effective when the distribution of fitness effects is narrow in range, i.e. of similar magnitude. Combining selection strengths of different magnitudes in the root mean squared method leads to more strongly deleterious mutations ($2Ns{\sim}100$) overshadowing the effects of very weakly deleterious mutations ($2Ns{\sim}1$). As a result, we show that our API implementation is expected to be highly accurate when users calculate $B'$ without a DFE (constant $2Ns$), and is also fairly accurate when only a single class of mutations is considered, though it is only somewhat accurate when modelling a DFE with mixed mutation classes. It is important to highlight that in the case of a DFE, while we do substantially underestimate $B'$, it is still a dramatically more effective estimate of diversity than $B$ calculated assuming no interference effects.

#### Caveats and recommendations

There are two major caveats of our $B'$ implementation in the API that remain, beyond modelling a DFE. First, we implement the framework of Becher and Charlesworth that models regions with no recombination but does not model low rates of recombination. Thus, our implementation provides no framework for modelling reduced recombination. Although HRI effects are typically maximized in non-recombining regions, low recombination regions with sufficient selected sites will also be impacted by HRI. We have not implemented the suggested methods in Text S4 of Good et al. (2014) to account for recombination and we encourage further research and development in this important area. Second is a concern important for users that may use the API to calculate $B'$ in a specific non-recombining region of the genome to adjust and improve a $B$-map. While such calculations may be accurate for the non-recombining region itself, this may not be true for the rest of the genome. This is because even when HRI is restricted to specific regions, HRI influences background selection genome-wide. To understand this best, consider a non-recombining region containing 10 kb of selected sites adjacent to a high recombination region. $B'$ can be accurately calculated for the non-recombining region experiencing HRI using the API, however there is no framework to accurately calculate the BGS effects that the 10 kb of selected sites in the non-recombining region inflict upon the high recombination region (Figure ST4.1). In the most extreme case, calculating $B$ assuming no interference effects for the site immediately adjacent to the HRI region will model all selected sites in the non-recombining region as being a recombinant distance of 1bp multiplied by the local recombination rate away, i.e. it will model 10 kb of selected sites as though they are all stacked on each other 1bp away, leading to $B{\sim}0$ (Figure ST4.1). In reality, the boundary is fuzzy between low-recombination regions experiencing increased HRI and higher recombination regions evolving under classic BGS, and HRI in one region decreases the contribution of those sites to BGS genome-wide (Figure ST4.1). To accurately capture the effects of regions evolving in an interference regime upon linked and unlinked sites outside an HRI region requires alternative methods of modelling background selection that explicitly account for interference effects (see Barroso and Ragsdale, 2025).

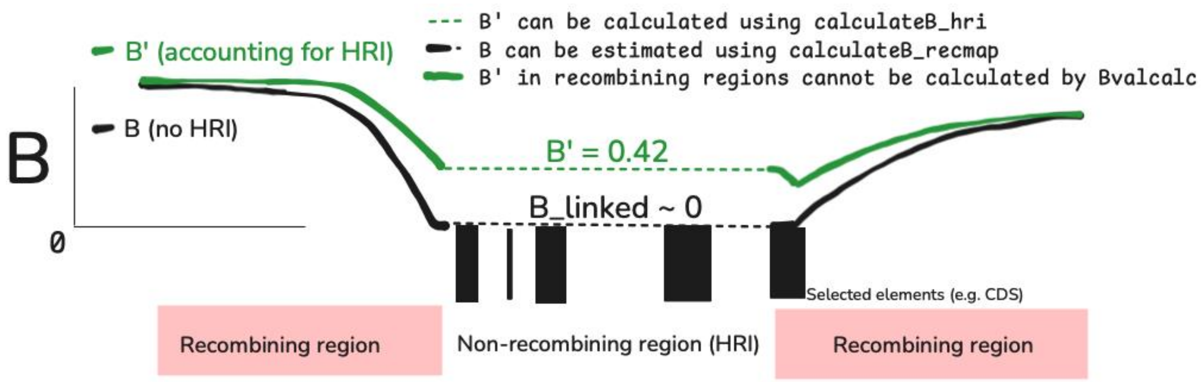


**Figure ST4.1. Illustrative diagram of the difference between $B'$ within and around a non-recombining region experiencing strong HRI effects (green lines that account for HRI), and $B$ assuming no HRI effects (black).** Note that the black lines are estimated by *Bvalcalc*'s primary modules by default, the dashed green line ($B'$ in a non-recombining region) can be estimated with the `calculateB_hri` API, but the solid green line ($B$' in a recombining region) cannot be estimated with *Bvalcalc*.

Despite the caveats with our API implementation for calculating $B'$, we hope it is useful to users working with non-recombining regions experiencing HRI and that it may aid theorists in population genetics to develop more comprehensive models of background selection. It may be useful to calculate $B'$ for some regions in genomes for which HRI effects are rare and isolated to improve a calculated $B$-map, though as our testing is limited, we advise users to use caution when applying the *Bvalcalc* $B'$ API in their research and recommend validating findings with appropriate simulations.
